# Compound-specific DNA adduct profiling with nanopore sequencing and IonStats

**DOI:** 10.1101/2025.10.31.685820

**Authors:** Yrjö Koski, Divyesh Patel, Natalia Kakko von Koch, Paula Jouhten, Lauri Aaltonen, Kimmo Palin, Biswajyoti Sahu, Esa Pitkänen

**Affiliations:** Institute for Molecular Medicine Finland (FIMM), University of Helsinki, Helsinki, Finland; Applied Tumor Genomics Research Program, Faculty of Medicine, University of Helsinki, Helsinki, Finland; The Norwegian Centre for Molecular Biosciences and Medicine (NCMBM), University of Oslo, Oslo, Norway; Department of Bioproducts and Biosystems, School of Chemical Engineering, Aalto University, Espoo, Finland; Department of Medical and Clinical Genetics, University of Helsinki, Helsinki, Finland; iCAN Digital Precision Cancer Medicine Flagship, Helsinki, Finland; Department of Cancer Genetics, Institute for Cancer Research, Oslo, Norway

**Keywords:** Nanopore sequencing, DNA adducts, genotoxicity, exposome, IonStats

## Abstract

Covalently bound DNA adducts are mutation precursors that contribute to aging and diseases such as cancer. Accurate detection of adducts in the genome will shed light on tumorigenesis. However, commonly used detection methods are unable to pinpoint the exact genomic locations of adducts. Long-read nanopore sequencing has the potential to accurately detect multiple types of DNA adducts at single-nucleotide precision. In this study, we developed a novel statistical toolkit, IonStats, to profile DNA adducts in nanopore sequencing data. With IonStats, we investigated the effects of the adduct-inducing genotoxic compounds aristolochic acid II and melphalan on nanopore sequencing and found both shared and compound-specific perturbations in base quality scores, ionic current profiles, and translocation dynamics. Notably, these treatments profoundly altered nanopore readouts and led to substantial sequence-specific read interruptions. Our study shows that nanopore sequencing can be effectively employed to detect and characterize DNA adducts, paving the way for high-resolution, high-throughput profiling of DNA damage and the exposome.

## 1 Introduction

DNA adducts are covalently bound lesions caused by exogenous and endogenous genotoxic compounds that react with DNA directly or after metabolic activation [1]. Enzymatic DNA repair mechanisms remove most adducts, but a small fraction may escape the repair [2]. Recent advances in DNA adductomics have revealed pronounced tissue- and age-specific heterogeneity [3, 4], while highlighting the influence of local chromatin and sequence context on adduct formation [5]. Adducts can interfere with DNA transcription and replication, and give rise to mutations that may lead to cancer [6]. Several compounds have been implicated in cancer initiation, including aristolochic acids (AAs), polycyclic aromatic hydrocarbons (PAHs), and aflatoxin B_1_ (AFB_1_), linked to urological, lung, and liver cancers, respectively [7, 8, 9, 10]. Beyond cancer, DNA adducts serve as molecular imprints of environmental exposures, contributing to accelerated biological aging and increased mortality [11]. Detecting and characterizing DNA adducts is thus crucial for understanding how exposure to DNA damage can lead to mutations and cellular dysfunction underlying both carcinogenesis and the progressive decline associated with aging.

There are two distinct classes of DNA adducts: monoadducts and crosslinks [12]. Monoadducts are formed through chemical modifications by small molecules on individual nucleotides. The primary targets of monoadduct binding depend on the reactive species, nucleophilicity of DNA sites, steric factors, and DNA sequence context. For small alkylating agents, the primary binding sites are the ring nitrogens N3 and N7, which are most nucleophilic in purine bases. For bulky aromatic compounds, such as AAs that form aristolactam (AL) adducts, the primary binding sites are exocyclic amino groups [13]. Molecules with multiple reactive groups can form DNA-DNA crosslinks, where two nucleotides are bound by the adduct [14]. Crosslinks can be either inter- or intrastrand, where the bound nucleotides are on different strands or the same strand, respectively [15, 16].

The most prominent methods for DNA adduct detection include mass spectrometry- and amplicon-based methods [17]. High-resolution mass spectrometry enables comprehensive profiling of adduct exposure, detecting most DNA monoadducts that are present in the sample [18, 19]. However, these methods require the DNA to be broken down into individual nucleotides, losing the positional information of the adducts in the genome [17]. Amplicon-based methods solve this issue by recognizing the adducts chemically or enzymatically by using DNA repair enzymes to mark and cleave DNA adducts [20, 21]. However, amplicon-based methods are unable to distinguish between different types of adducts [17].

Nanopore sequencing is an emerging technology with the potential to differentiate between adduct types at single-nucleotide resolution while preserving site-specificity [22, 23]. Nanopore sequencing measures the ionic current as a single-stranded DNA fragment translocates through a nanopore, which is embedded in an electrically insulating membrane. The nucleobases that reside inside the nanopore at any given time alter the ionic current based on their physical structure. Hence, analysis of the ionic current can potentially reveal both the DNA sequence and modifications such as adducts [24, 25]. For instance, in SMAdd-seq, DNA adducts are induced at linker DNA with angelicin, generating distinct nanopore signatures that enable chromatin accessibility profiling [26]

Most previous studies have focused on adducts only in specific known sequence contexts [22, 23]. In this study, we show that nanopore sequencing can be used to characterize adducts in arbitrary sequence contexts. We implemented a novel software package called IonStats for distilling nanopore sequencing data into multiple statistics revealing deviations from the unadducted state. We show that these statistics are informative for adduct formation by creating and analyzing a novel nanopore sequencing dataset of whole-genome amplified *Saccharomyces cerevisiae* DNA with two adduct-inducing genotoxic compounds: aristolochic acid II (AAII) and melphalan. We conducted experiments on cisplatin and mitomycin C (MMC) as well, but these samples were later excluded from the analyses due to data quality issues (Supplementary methods). We demonstrate distinct treatment effects by comparing the sequencing statistic distributions between treated and untreated samples. We also observed that some treatments increase read interruptions, where DNA fragments are not fully sequenced, likely due to bulky adducts or crosslinks bound at specific sequence contexts. IonStats is available open source at GitHub (https://github.com/ykoski/ionstats).

## 2 Materials and Methods

### 2.1 Whole genome DNA amplification

Yeast genomic DNA was extracted from the haploid *S. cerevisiae* strain CEN.PK113-7D by using the MasterPure Yeast DNA Purification Kit (MPY80200, LGC). High molecular weight yeast genomic DNA was uniformly amplified by using the REPLI-G mini whole genome amplification kit (150023, Qiagen). In brief, 20 ng of yeast genomic DNA was used as a template and amplified by incubating with REPLI-G DNA Polymerase at 30 °C for 16 hours. Amplified yeast genomic DNA was purified by using 1.8x Ampure XP beads (10136224, Beckman Coulter) and eluted in 10 mM Tris-HCl, pH 8.0, or double-distilled water. Amplified yeast genomic DNA was fragmented and size-selected for 10–15 Kb by using g-tube (520079, Covaris). For the melphalan treatment, DNA was extracted with ultrapure distilled water (10977035, Invitrogen).

### 2.2 Aristolochic acid II treatment

AAII DNA adducts were formed by using the protocol described in [27]. Amplified yeast genomic DNA (10 µg DNA, one µg per µl) was treated with 0.25 mM AAII solution in 50 µl of 50 mM potassium phosphate buffer, pH 5.8. AAII was activated with 1 mg of zinc dust and incubated at 37 °C for 30 minutes. AAII-treated DNA was purified by using 1.8x Ampure XP beads and eluted in 25 µl 10 mM Tris-HCl, pH 8.0.

### 2.3 Melphalan treatment

Melphalan DNA adduct was prepared by using the protocol in [28] with a modification. Amplified yeast genomic DNA was eluted in distilled water (10 µg in 10 µl water). Amplified yeast genomic DNA was incubated with 8 µl of melphalan stock solution (1 mg per ml of DMSO) and 12 µl of DMSO at 37 °C for 1 hour. Melphalan-treated DNA was purified using Phenol-Chloroform extraction and eluted in distilled water.

### 2.4 DNA sequencing and data collection

Two replicates for each treatment were prepared for sequencing. Approximately 400 ng of DNA was used for each sample to prepare the sequencing library with the Native Barcoding Kit 24 V14 (SQK-NBD114.24, Oxford Nanopore Technologies (ONT)) following the manufacturer’s protocol. In total, 13 barcodes were used, taking into account the discarded samples (two for each treatment, two for controls, and one additional for cisplatin size-selected; Supplementary methods). The constructed libraries were loaded onto the R10.4.1/FLO-PRO114M flow cell on the PromethION 24 device. The sequencing and data collection were performed with the MinKNOW software v24.02.19, which generated .pod5 files containing the raw ionic current measurements for individual DNA molecules.

### 2.5 Data processing

Basecalling and alignment were performed with ONT’s basecaller Dorado (version 0.6.0) with the dna_r10.4.1_e8.2_400bps_sup@v4.3.0 model. The aligner function in Dorado uses minimap2 [29] to align the basecalled reads to a reference genome with the “lr:hq” preset. We used the *S. cerevisiae* strain CEN.PK113-7D assembly from [30] as the reference genome. To map the raw ionic current measurements to positions in the reference genome, a signal-alignment step is required. The ionic current is divided into events, and each event is mapped to a reference position. We applied Uncalled4 [31] to align ionic current measurements to reference positions for all reads. This allowed us to compute and compare ionic current statistics for different samples.

Uncalled4 normalizes each read with a linear transformation fitted to the pore model, matching the read’s event mean currents to the expected currents of its aligned *k*-mers and refining the fit by linear regression ([31], Methods). Both steps estimate the read’s scale from a non-robust statistic, so a read carrying a small population of low current events has an inflated scale estimate that compresses every other event toward the read center, and pooling across reads turns this into an apparent level-dependent shift. For this reason, we used the current means collected by Uncalled4 to normalize the current means and current SDs using a robust Interquartile Range (IQR) normalization

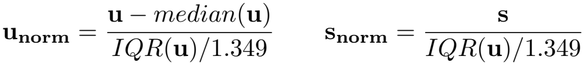

where **u** are the current means and **s** are the current SDs of a read. The IQR was divided by 1.349 = Φ^*−*1^(0.75) *−* Φ^*−*1^(0.25), which is a normal-consistency constant of the IQR range, so that the denominator estimates the standard deviation for Gaussian data.

### 2.6 Modkit validation dataset

We downloaded a dataset described in [32] to validate and benchmark IonStats on a modification detection task on three methylation types. The modkit validation dataset consists of short synthetic oligos that cover 5mC, 5hmC, and 6mA methylations in all 5-mer contexts. The reference genome consists of 32 short contigs (155 bp each) carrying methylations at specific positions (8 sites per contig, 256 in total). The set consists of two replicates per treatment along with two untreated control samples. We used only a subset of data predefined by the dataset authors [32], which contains 5000 randomly selected reads per synthetic construct per sample. The samples were preprocessed with the same pipeline as the adduct-treatment dataset.

### 2.7 Data analysis of sequencing statistics with IonStats

IonStats can extract and compare quality scores, ionic current means and standard deviations, dwell times, and adjusted dwell times between treated and control samples.

The quality score (*Q*) is a value given to each base by the basecaller, and it is logarithmically related to the estimated basecalling error probability (*P*): *Q* = *−*10 log_10_ *P* . The ionic current mean and standard deviation are computed from the raw ionic current measurements that belong to a single event, defined by the signal alignment. The dwell time refers to the time that a *k*-mer resides inside the nanopore, and we quantified it as the number of measurements that belong to an event. To capture the interaction between putative adducts and the nanopore motor protein, we computed the motor-protein-shifted dwell times (adjusted dwell times), where the dwell time of a *k*-mer was associated with a *k*-mer 12 base pairs downstream in the reference genome. Reported estimates of the motor protein offset span roughly 11-14 bases [31, 23].

IonStats collects values for selected variables and groups them by the reference *k*-mer or by reference position for a specific chromosome or contig. It then performs distributional tests on the *k*-mer and reference position distributions to find the *k*-mers or positions that show statistically significant differences in the variables of interest. It applies the Kolmogorov–Smirnov (KS) test, the Cramér–von Mises (CvM) criterion, and the Anderson-Darling (AD) test to compare the *k*-mers. For all of these tests, we used the versions that were implemented in the SciPy [33] Python library. These three tests are applied to capture different effects since KS and CvM are more sensitive to differences in medians, while AD is more sensitive to differences in distribution tails. The KS and CvM tests assume continuous data, which the quality score and dwell times violate. For AD, we applied a tie-corrected midrank variant. Adding a uniform tie-breaking noise to the discrete statistic had little effect (Suppl. Table 14, 15), so the results are reported without it. To control for multiple testing, IonStats applies the Benjamini-Hochberg correction for the *p*-values of each test with a significance level *α*. We used *α* = 0.05.

### 2.8 Quantile difference analysis

To study how the extreme values of *k*-mer distributions compare in treated and control samples, we computed the top and bottom 1%-quantiles for each sequencing variable. We then computed the quantile difference to measure if a *k*-mer shows more extreme values in the treated samples. The quantile difference for *k*-mer *i* is defined as: *d*_*i*_ = *q*_*t,i*_ *− q*_*c,i*_, where *c* is control and *t* is treatment, and *q* can be either the lowest or highest 1%.

### 2.9 Data analysis for interrupted reads

Nanopore sequencing reads can exhibit low rear barcode quality, indicating that the read barcode was non-determinable based on the sequence at the end of the read. This led us to believe that interrupted reads, *i*.*e*., reads that interrupted the sequencing and did not pass completely through the pore, belong to this subset of reads. We classified reads as interrupted based on the `barcode_rear_score` value, which is recorded for all reads in the sequencing summary file provided by MinKNOW and which ranges between *−*100 and 100. If `barcode_rear_score` was *<* 60, we classified the read as interrupted. We chose this threshold as MinKNOW requires a minimum value of 60 from either the front or rear barcode scores for a read to be assigned to a barcode.

After further investigation, we noticed that some reads classified as interrupted had a low ionic current at the end of the read, while other reads had a similar ionic current level at the end of the read to fully-read reads (Suppl. Fig. 4). We divided the interrupted reads into two classes based on their last 20 ionic current measurements: passing-like reads with high end-mean, and blocking reads with low end-mean (*<* 50 pA). The passing-like reads exhibit a similar current level at the end of sequencing as fully-read reads, and the blocking reads exhibit a lower current level. We analyzed the end positions of these reads by extracting 25 bases upstream and downstream (51 in total) of the mapping end position in the reference genome to study whether the reads typically ended at specific sequence motifs, which might be binding sites of adducts. We also performed a motif discovery analysis on subsequences at the mapping end position and on sequences that interact with the motor protein.

### 2.10 Motif discovery analysis

To search for patterns that could correspond to recurrent adduct-forming sequence motifs in the DNA, we performed motif discovery analysis with STREME [34] for the top-5000 *k*-mers ranked by the AD-test value. In the STREME analyses, we used a coverage-matched set of 5000 *k*-mers as the control sequences, with no overlap with the treated set. To process the *k*-mers only in forward-strand orientation, we used a custom STREME alphabet, which did not consider the complementarity of nucleobases. The motif discovery was performed separately on two independent replicates. A motif had to occur in both replicates with an *E*-value smaller than 0.05 to be reported in Fig. 5 and considered significant. We used the Logomaker [35] Python library to draw sequence logos of the interrupted read position-weight matrices in Fig. 6.

Motifs discovered in the two replicates were compared with TOMTOM [36], which measures the similarity of two position weight matrices. A pair was counted as replicating only when both motifs independently cleared the STREME significance cut (*E <* 0.05) in their own replicate. In addition, we tested motifs between treatments to assess whether a motif was treatment-specific.

### 2.11 Modification detection from test scores

The distributional two-sample tests are performed at every reference position for each of the four statistics. Significant positions are called by a decoy threshold, given by a control-control comparison that provides a null distribution for each statistic without modifications. The threshold is the most permissive value at which the counted false discovery rate (FDR), null positions above the threshold divided by observed positions above it, stays within the target (decoy FDR set to 0.01 by default). If a second control sample is not available, the method will resolve to a Benjamini-Hochberg FDR correction at *α* = 0.05.

To limit the number of false positive calls, and to improve the localization of the modification calls, we apply a two-step process: i) Peak detection: local maxima in the test scores of the significant positions are marked, and the maximum within a window of *±sep −* 1 positions is selected (*sep* = 2 by default). Here *sep* refers to the minimum separation of called positions, *i*.*e*., separation of 2 means that calls can be made in every second position. ii) Snapping: Each peak is moved to the nearest candidate base within a *±sep −* 1 window, if the candidate base is known. If there are no candidate bases available in the window, the peak is discarded. If the candidate base is not known, instead of snapping, peak merging is applied. Surviving peaks are merged: a peak is retained if no other peak within *±merge* carries a larger statistic (*merge* = 5 by default). In order to meaningfully merge peaks, *merge* ≥ *sep* must hold.

In our validation experiment, we used a decoy FDR value of 0.002 and a peak detection *sep* = 7 for the modkit validation dataset, after conducting a hyperparameter search on *sep* values (Suppl. Table 4). For scoring, we used a tolerance of one, meaning that a call is a true positive if it lands within *±*1 bases of the true site.

Modification detection is only applicable in the reference position mode, as the pipeline combines information from neighboring positions, consistent with the hypothesis that a modification perturbs many consecutive events.

### 2.12 Positional regression analysis

We applied a positional regression analysis to quantify which base, at which position relative to the sensed base, is associated with a larger adduct-induced shift of the low tail of the current mean distribution. For each *k*-mer with at least 500 observations, we used the lowest 1% quantile value *q* to define the shift response for each *k*-mer *k*:

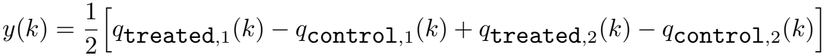

where each treatment replicate was paired with a different control replicate. We stratified the analysis by the sensed base at index 6 (*i*.*e*., the seventh base of the nine-base sensing window, with most influence on the current level in the R10.4.1 9-mer model [31]), and fit an ordinary least-squares model for each sensed base. The regression model contains 24 indicator variables, one for each non-A base for each position after the sensed base stratification, and an intercept variable.

To make the results more interpretable, the coefficients are assembled into a 4 *×* 9 matrix *M* over base and position, and each position column is centered to mean zero. A displayed value in the heatmap Figure 5 is the effect of that base relative to the average base at that offset.

To indicate which coefficients reproduce between replicates, we mark them against a standard error estimated from the replicates themselves. The two treated replicates are additionally fit separately, one against each control replicate, giving *M*_1_ and *M*_2_. The replicate standard error is the pooled standard deviation of the half-differences (*M*_1_ *− M*_2_)*/*2 across all cells, sensed bases, and treatments. We opted for a replicate-based standard error, as the model-based standard errors proved to be overconfident, as they describe sampling within a single replicate, and we observed the between-replicate error to be much larger.

### 2.13 Analysis of ionic current drop events

A current drop was defined as a decrease in ionic current per event relative to the expected level of its *k*-mer. For event *i* assigned to *k*-mer *j*, we computed the distance *r*_*i*_ = *u*_*i*_ − *µ*_*j*_, where *u*_*i*_ is the event’s normalized mean current and *µ*_*j*_ is the median of that *k*-mer’s events in the control sample, and classified the event as a current drop when *r*_*i*_ *< τ* . We used *τ* = *−*1.21. The threshold is a fixed operating point, and the enrichment of low current events is monotone with *τ*, while the control-control comparison remains within 5% of unity across the range −0.8 to −2.0 (Suppl. Table 13). Current drops were restricted to sustained low events using *l*_*i*_ = *log*_10_(*s*_*i*_*/σ*_*j*_), the log-ratio of the event’s current standard deviation *s*_*i*_ to that *k*-mer’s control value *σ*_*j*_. Events with *l*_*i*_ ≤ 0.6 were kept as sustained drops, and those with *l*_*i*_ *>* 0.6 were discarded as they have elevated within-event variance and represent transitions between levels rather than a stable displacement.

Consecutive current drops were measured in a window of nine events centered on each current drop, the span of a single 9-mer. Each event in the window was compared against its own expected level, and an event was counted as part of a run when its distance fell below *τ/*2. The run length is the number of contiguous low events containing the central drop.

## 3 Results

### 3.1 IonStats: a comprehensive toolkit for capturing adduct effects in nanopore sequencing data

Nanopore sequencing generates large volumes of ionic current data, posing substantial challenges for downstream analyses. To distill informative features for adduct detection, we developed a computational workflow called IonStats. IonStats collects and summarizes diverse statistics from nanopore current and basecalled sequencing data by *k*-mers or reference genome positions. Comparison of the statistics of treated samples with those of untreated samples allows the characterization of treatment effects. In addition, IonStats collects read-level statistics including read counts and lengths, and base quality scores. In our study, we used *k* = 9, matching the number of nucleotides that reside within the nanopore (ONT pore version R10.4.1) at any given time [31]. This choice also ensured high *k*-mer coverage in our experiments.

For each *k*-mer or reference position, IonStats collects nanopore current means and standard deviations, base quality scores, and dwell times. The latter indicates the number of measurements obtained for each nucleotide, correlating with the time the nucleotide resided in the nanopore sensing region. In addition, IonStats computes motor-protein-adjusted dwell time (ADT). This statistic reflects the duration that each nucleotide resides at the motor protein, rather than within the sensing region. ADT lets us study the effects that adducts might have when interacting with the motor protein, in addition to in-pore effects. Finally, adducts may cause failed translocations, which IonStats can detect by identifying interrupted reads through missing rear barcode sequences.

IonStats provides distributional tests to detect differences between sample groups, typically between treated and control samples. The available tests include the Kolmogorov-Smirnov (KS) test, the Cramér-von Mises (CvM) criterion, and the Anderson-Darling (AD) test. Compared to the more common KS test, AD is more sensitive to differences in tails of distributions. Together, these complementary tests enable robust detection of treatment-induced perturbations in nanopore data.

### 3.2 IonStats outperforms ELIGOS in methylation detection

We applied IonStats to the ONT modkit validation dataset [32] to evaluate its performance in a modification detection task. We benchmarked IonStats against ELIGOS [37], which uses the percent error of specific bases (%ESB) metric to detect DNA or RNA modifications without modification-specific training. ELIGOS was evaluated with three different modification detection pipelines: the same as IonStats (Matched pipeline), *sep*-optimized pipeline (Best separation), and the tool’s own calling rule (ELIGOS calling rule). Our benchmarking showed that IonStats outperformed ELIGOS for all three methylation types in the dataset (Table 1, Suppl. Figure 3) on all current-derived statistics. The only IonStats statistic that showed worse performance than ELIGOS on two of the three methylations was the quality score, which is not derived from the raw ionic current. The best-performing IonStats statistics were the current mean and current SD with average *F*_1_-scores of 72.6 and 72.4, respectively. While current mean showed better performance on cytosine methylations by 9 to 10 percentage points, current SD outperformed current mean on 6mA by 19 points. In this comparison, the candidate methylated bases were given, and snapping was enabled (Methods). When candidate bases were not given, performance remained approximately at the same level, although the localization of methylations worsened (Suppl. Table 2).

**Table 1:** IonStats recovers more modified sites at higher precision than ELIGOS for all three modifications with the current-derived values. Recall (Rec.), precision (Prec.), and percentage *F*_1_, on the modkit validation dataset of 256 modified sites at full depth (166,000 reads, 4850*×* median coverage, 100% modified). All entries use the two-sample KS statistic; the CvM and AD statistics perform equivalently on this panel, within the variation between replicates (Suppl. Table 1). IonStats was run through a calling pipeline with: a control-versus-control decoy threshold, peak separation 7, snapping to the nearest candidate base within 6 bp, and a positional tolerance of *±*1 bp against the same truth set. For ELIGOS, we applied the exact same pipeline, the same pipeline with an *F*_1_-optimized peak separation value of 2, and the tool’s own calling rule. Each entry is the mean of two independent replicates. The right-hand column is the mean of the three *F*_1_ values, and the best *F*_1_ per modification is in bold. The basecall quality score is the basecall-level baseline: it uses no ionic current information and is reported for reference.

|  | 5mC |  |  | 5hmC |  |  | 6mA |  |  |  |
|---|---|---|---|---|---|---|---|---|---|---|
| Statistic | Rec. | Prec. | *F*<sub>1</sub> | Rec. | Prec. | *F*<sub>1</sub> | Rec. | Prec. | *F*<sub>1</sub> | Mean *F*<sub>1</sub> |
| IonStats |  |  |  |  |  |  |  |  |  |  |
| Current mean | 91.8 | 84.2 | **87.9** | 87.7 | 78.1 | **82.6** | 50.6 | 44.7 | 47.4 | **72.6** |
| Current SD | 76.2 | 78.3 | 77.2 | 74.8 | 72.4 | 73.6 | 63.5 | 69.9 | **66.5** | 72.4 |
| Dwell time | 66.0 | 70.9 | 68.4 | 64.8 | 60.6 | 62.6 | 45.3 | 50.0 | 47.5 | 59.5 |
| Quality score | 33.4 | 48.2 | 39.4 | 36.3 | 49.9 | 42.0 | 32.6 | 46.1 | 38.2 | 39.9 |
| ELIGOS |  |  |  |  |  |  |  |  |  |  |
| Matched pipeline | 49.0 | 46.2 | 47.6 | 45.9 | 41.0 | 43.3 | 29.1 | 29.3 | 29.2 | 40.0 |
| Best separation | 67.8 | 37.7 | 48.4 | 74.2 | 36.0 | 48.5 | 59.0 | 27.7 | 37.7 | 44.9 |
| ELIGOS calling rule | 76.4 | 29.0 | 42.0 | 88.5 | 26.3 | 40.5 | 66.6 | 19.4 | 30.0 | 37.5 |

Out of the three methylations, 6mA was the most difficult to detect. We visualized the effect that each methylation type induces on different variables by pooling the observed differences across all modified positions between treated and control samples (Figure 2A). We observed that the localization of 6mA was the most challenging, as the induced effect of the modification is shifted from the sensed position. The highest peak in the current mean difference against unmodified DNA is positive for 6mA, while it is negative and well-localized for the cytosine methylations. However, the standard deviation effect is better localized in 6mA compared to the cytosine methylations, even though the magnitude of the peak is smaller.

Next, we studied the effect of modification fraction and coverage on modification detection (Figure 2B,C). As expected, IonStats performance declined as the modification fraction decreases. However, at 50% modifications, the *F*_1_-score was better than in fully modified samples for current mean in 5mC and 6mA. The recall remained at a similar level, while the precision improved, resulting in an increased *F*_1_-score (Suppl. Table 3; precision odds ratio = 2.0, *p* = 1.6 *×* 10^*−*4^).

The coverage had little effect on the *F*_1_-score for ionic current mean. However, for ionic current SD and for ELIGOS detection, the performance worsened as the coverage decreased.

### 3.3 Adduct-induced DNA damage reduces nanopore read length and quality

We first examined whether the effects of adduct treatments can be observed in read-level nanopore sequencing statistics after treating *S. cerevisiae* DNA with two genotoxic, adduct-inducing compounds (Fig. 1A; Materials and Methods). These statistics included the number of reads sequenced (Fig. 1B), read lengths and mean base quality scores (Fig. 1C), and rear barcode quality scores (Suppl. Fig. 1, Suppl. Table 5). Both AAII and melphalan samples showed lower mean quality scores (16.4 and 15.2, respectively; Suppl. Table 5, 7) than untreated controls (19.6, *p <* 10^*−*16^ for all compounds) (Fig. 1C). Melphalan samples had shorter reads (median lengths: 1833) compared to AAII and control samples (AAII 6744, control 7061; Suppl. Table 5, 7). The replicates generally correlated very well, except for AAII, where the two replicates differed in mean quality score (15.8 and 17.1; *p <* 10^*−*16^).

**Figure 1:**
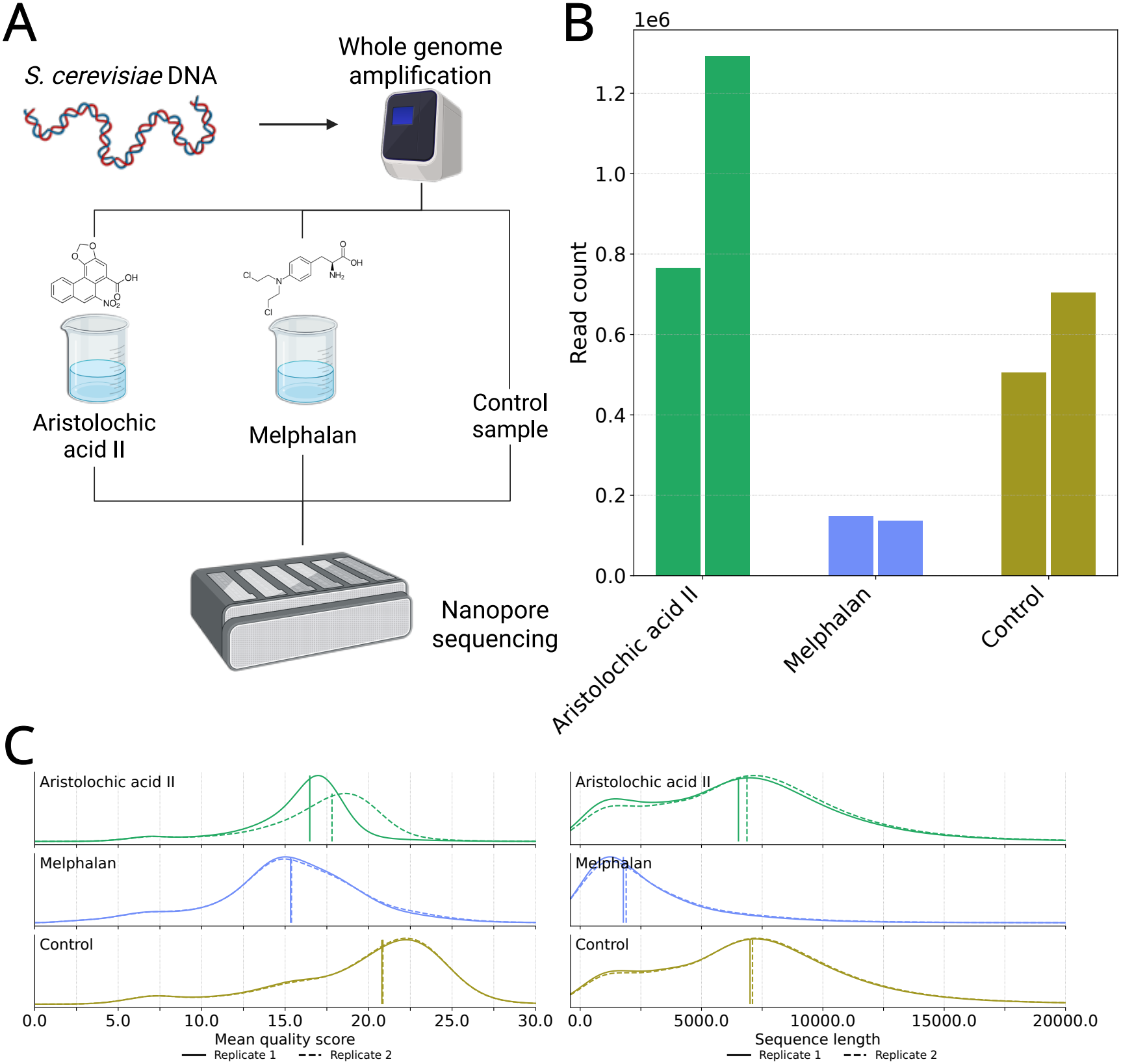
Overview of the experimental setup and sequencing statistics in the nanopore sequencing dataset. **A** Visualization of the experimental setup of this study. **B:** Number of reads per sample replicate obtained in the experiment. **C:** Read mean quality scores and read lengths by sample. The vertical line shows the median value of each replicate.

**Figure 2:**
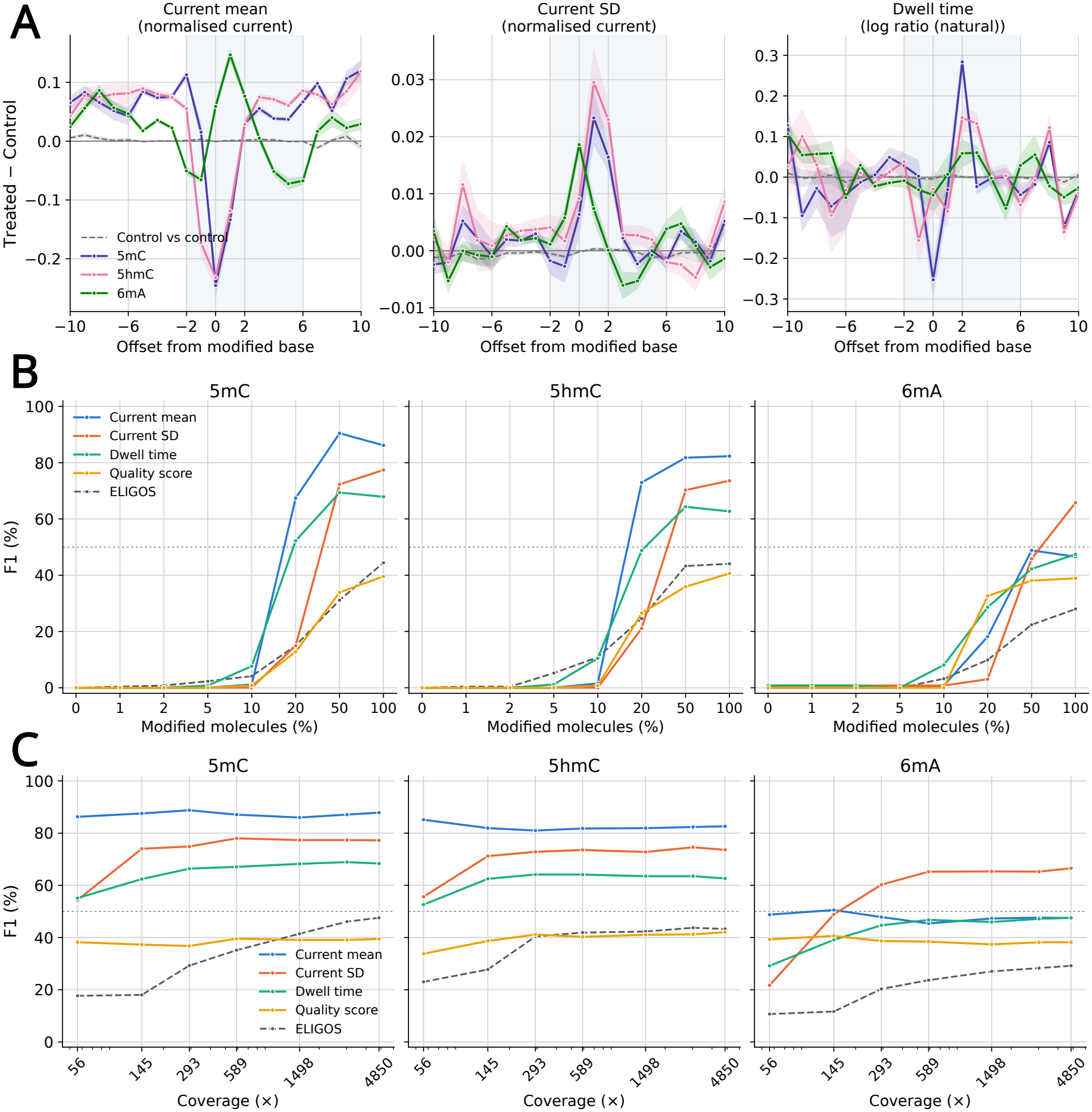
IonStats and ELIGOS modification detection benchmarking on methylation panel dataset. **A:** Modification profiles for 5mC, 5hmC, and 6mA for ionic current mean and standard deviation, and dwell time. The profiles show the averaged effect of each methylation type across all modified positions in the methylation panel dataset. The envelope shows the 95% confidence intervals. **B:** *F*_1_-scores for IonStats and ELIGOS modification detection for all three methylation types for varying modification fractions in the methylation panel dataset. The IonStats results are shown for all four data types. The methylated and control reads were mixed at different ratios to produce the samples with varying methylation fractions. **C:** *F*_1_-scores for IonStats and ELIGOS modification detection with different coverages for different methylations.

### 3.4 Treatment effects are observed in tails of nanopore current-level statistics

We then focused on the current-level statistics obtained with IonStats. Overall, for most treatments, the distributions of these *k*-mer-level statistics were consistent with those observed in controls (Suppl. Table 8 & 9, Suppl. Fig. 5-12). However, for AAII, melphalan, and cisplatin, the variability of many statistics differed substantially from controls (Suppl. Table 8 & 10, Suppl. Fig. 5-12). This led us to believe that the treatment effects are primarily observed at the tails of *k*-mer value distributions.

To directly investigate the low-current events, we analyzed the raw ionic currents to find events that have a much lower ionic current mean than expected (Methods). These are referred to as current drop events. Examples of current drop events in AAII, melphalan, and control samples are shown in Figure 3A. In these events, the ionic current mean was at a lower level compared to the median value of the control current mean.

**Figure 3:**
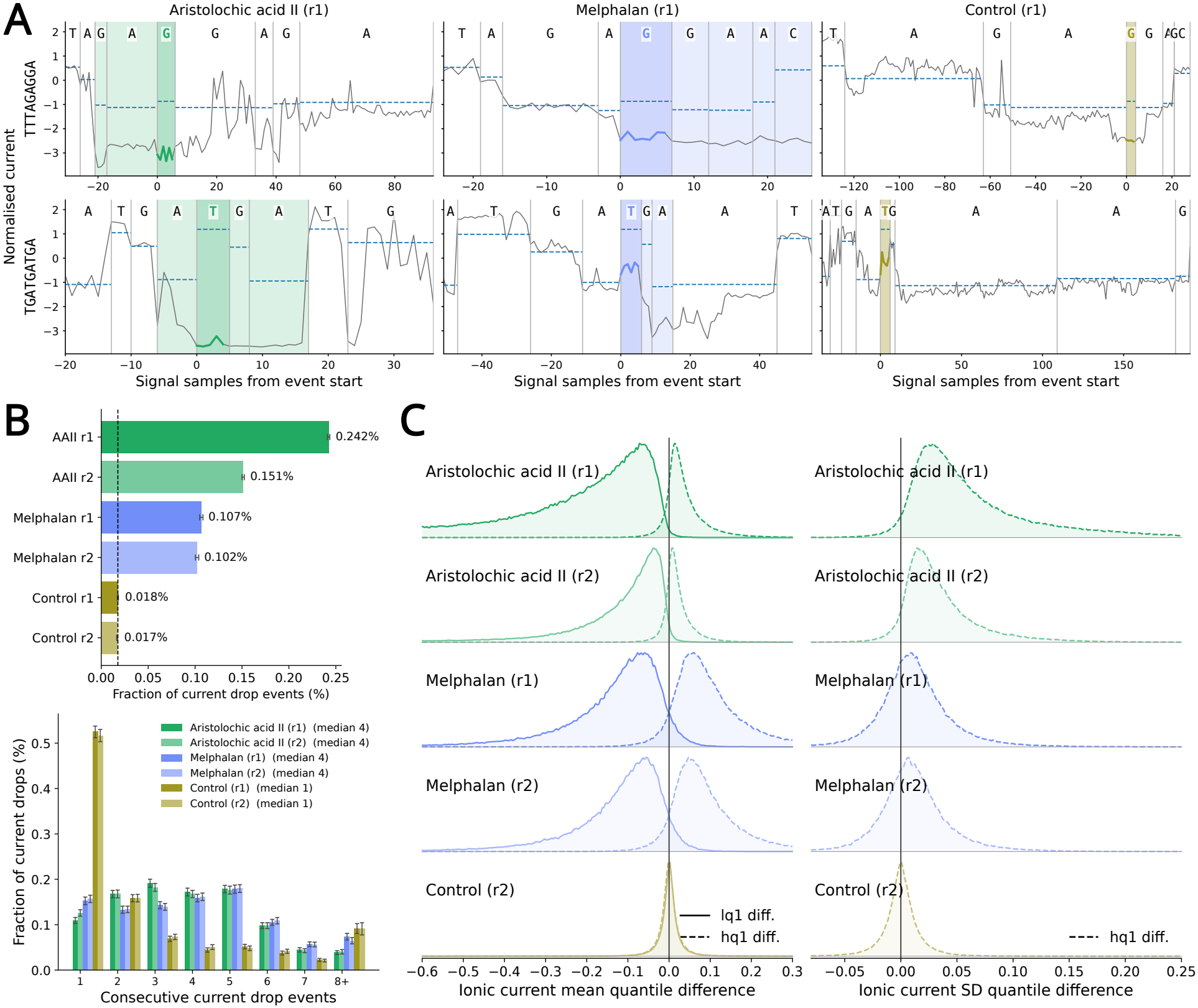
**A:** Examples of current drop events in AAII, melphalan, and control samples for two matched *k*-mers. The darker shading highlights the current drop event at that *k*-mer, the lighter shade highlights the surrounding current drop events, the blue dashed line shows the control median ionic current level for the *k*-mer, and the letters indicate the sensed base at every event. **B:** The top panel shows fractions of current drop events out of all events in all AAII, melphalan, and control samples. The bottom panel shows the frequency of consecutive current drop events in each sample. **C:** Ionic current mean and SD quantile difference densities in AAII, melphalan, and control (r2) samples compared against control (r1).

Current drop events were 6-to 13-fold more frequent in the AAII and melphalan samples than in the controls (Figure 3B). Additionally, the current drops typically spanned many consecutive events in the AAII and melphalan samples, with a median drop length of 4 events, compared to controls, which showed mostly singular drop events. This result suggests that both AAII and melphalan treatments induce current drops that span multiple events, that are likely caused by a bulky monoadduct translocating through the nanopore and disrupting the ion flow.

Consistent with this hypothesis, the extreme quantiles (1st and 99th percentiles) of ionic current statistics revealed pronounced shifts (Fig. 3C, Suppl. Table 11 & 12), while their means showed only minor differences (Suppl. Table 8 & 9). Specifically, the current mean low quantiles for most *k*-mers in AAII and melphalan samples were lower than those in controls (Fig. 3C). Furthermore, many *k*-mers displayed markedly higher variability in current levels, particularly in AAII samples. These observations support the notion that adduct formation affects only a subset of DNA nucleotides, producing a mixture of modified and unmodified *k*-mers (Fig. 3 A, B). Due to insufficient *k*-mer coverage, cisplatin samples were excluded from these analyses (Suppl. Table 5).

### 3.5 Adduct treatment induces more extreme dwell times

We noticed that the frequency of current drop events depended on the dwell time (Figure 4A,B). In AAII, melphalan, and control samples, the current drops occurred primarily in short events, with a dwell time less than 20. However, we observed that the current drop events tended to be longer in the AAII and melphalan samples compared to the controls, as their ratio with one control replicate increases steadily from the lowest dwell times up to around 30, while the other control replicate’s ratio stays near one.

**Figure 4:**
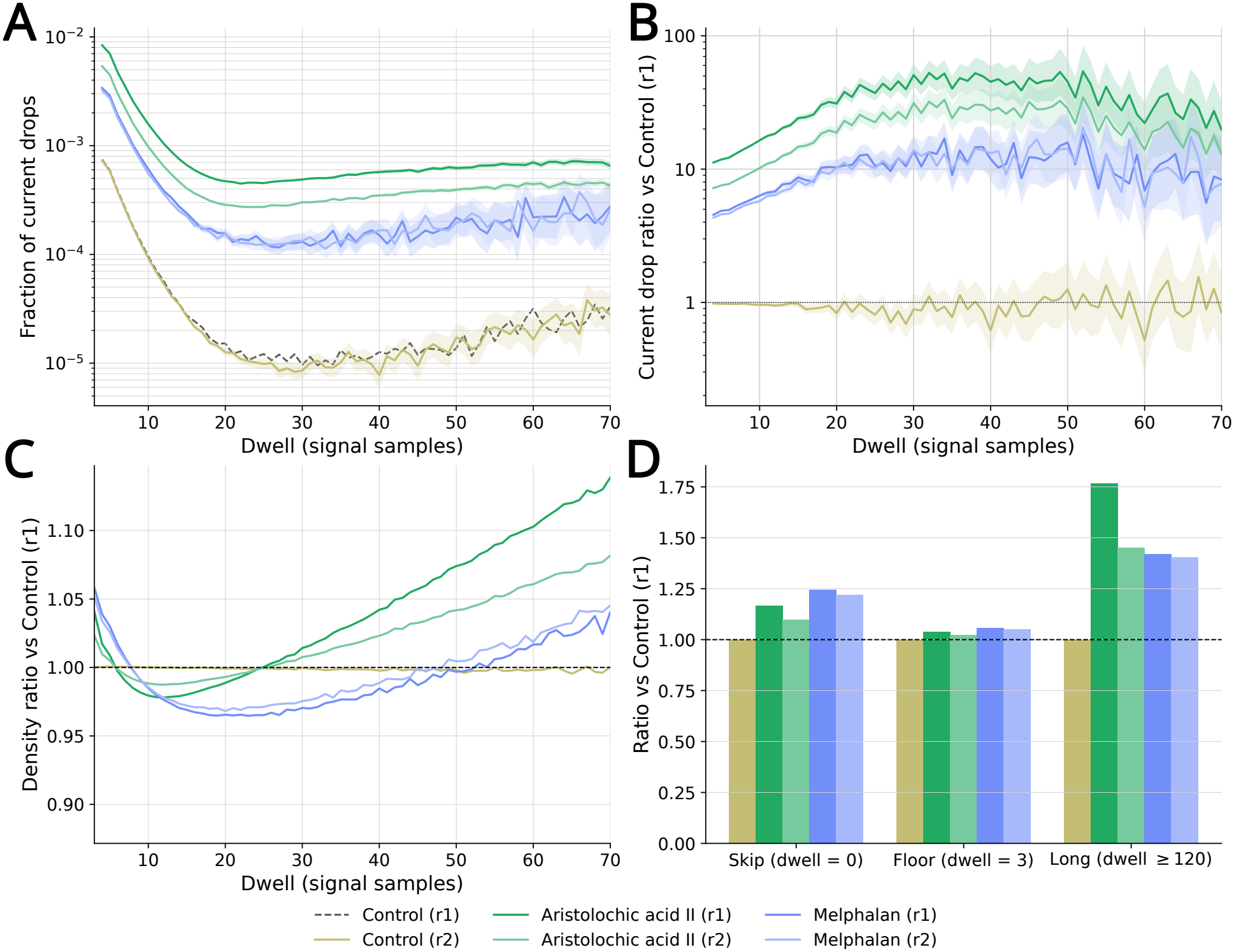
**A:** Frequency of current drop events as a function of dwell time in AAII, melphalan, and control samples. **B:** The current drop rate versus the control (r1) sample as a function of dwell time for AAII, melphalan, and control (r2) samples. **C:** The ratio of dwell time densities of all events versus control (r1) for AAII, melphalan, and control (r2) samples. **D:** The ratio of skipped events, shortest aligned events, and long-dwell events versus control (r1) in AAII, melphalan, and control (r2) samples.

After looking into the dwell times of current drop events, we wanted to compare dwell times between treatments in all events. We observed that there are more short dwell times, fewer medium-length dwell times, and again more long dwell times in AAII and melphalan samples (Figure 4C). While the trend was similar in both treatments, the range of decreased medium-dwell times was different. We then checked the ratios of specific dwell time classes: skipped *k*-mers (*i*.*e*., Uncalled4 did not align any current values to the *k*-mer), floor length of three, which is the shortest aligned event that Uncalled4 produces, and long dwells, with more than 120 measurements. We observed that there were more skip events in both treatments (*k*-mer-matched Student’s t *p <* 10^*−*16^ for AAII and melphalan). Additionally, there was a large enrichment of long dwell times (*k*-mer-matched Student’s t *p <* 10^*−*16^ for AAII and melphalan; Figure 4D).

These findings suggest that the current drops were most often associated with short events, while the longer current drop events were enriched in AAII and melphalan samples compared to the controls. Both treatments changed the dwell time densities across all events, as they showed more short and long events. We believe that the long events were caused by motor protein stalling due to interaction with bulky monoadducts. The surplus of low dwell times would suggest that the treatment increased the overall translocation speed. However, this could also be a current-alignment artifact, as adducted *k*-mers’ current level can vary a lot from the expected level, as we previously quantified.

#### 3.6 Treatment effects are driven by high purine content

In order to assess the sequence specificity of the treatment effects, we conducted a positional regression analysis to predict the difference in the lowest 1% of current mean values for all *k*-mers in AAII, melphalan, and control (Materials and Methods). The analysis was stratified by the sensed base, and the coefficients are shown in Figure 5. Drops in AAII are associated with VGAS at sensed A, CGCR at sensed C, VRGA at sensed G, and VATR at sensed T (IUPAC notation, offsets -2 to 1). Drops in melphalan are primarily associated with G-rich contexts such as NGAG at sensed A, GGCG at sensed C, GRGG at sensed G, and GRTG at sensed T. There was no sequence specificity in the low-quantile differences between two control samples, as no base-position pairs exceeded the replicate-based standard error threshold, and with *R*^2^ ≤ 0.003 against treated 0.22–0.52. This highlights that the observed effects in treated samples are treatment-specific.

**Figure 5:**
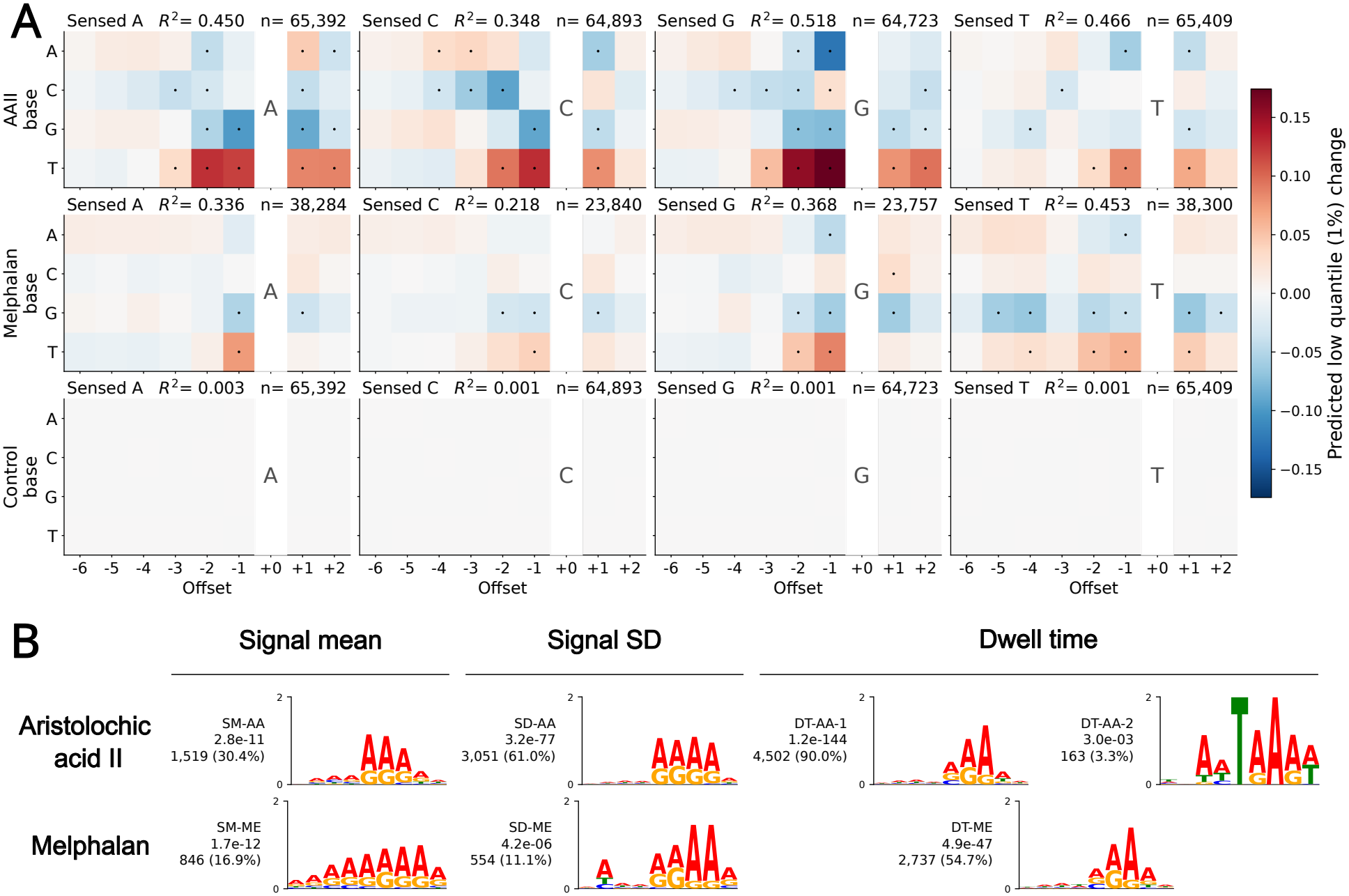
**A:** Positional regression coefficients predictive of a change in the lowest 1% quantile of ionic current mean values stratified by sensing base in AAII, melphalan, and control samples. Black dots highlight base-position interactions that reproduce between replicates after replicate-based standard error thresholding (Methods). Cells whose coefficient exceeded twice the standard error were marked with a black dot. **B:** All significant sequence motifs associated with the current mean and standard deviation, and dwell time, with motif discovery analysis on the top-5000 *k*-mers based on the AD test statistic. Next to each motif is its identifier, E-value, and the number and fraction of occurrences in the motif set. A motif had to be found in both replicates independently to be reported.

Additionally, we asked whether treatment-driven effects might be linked to recurring DNA sequence contexts. To explore this, we conducted sequence motif discovery on the top-5000 *k*-mers ranked by the test value, associated with differences in sequencing statistics between the treated and control samples, as determined by the AD-test (Methods). This analysis revealed one significant motif for current mean and current SD in melphalan and AAII, and two significant motifs in dwell time in AAII and one significant motif in melphalan dwell time (Figure 5B) with low *E*-values (*E <* 0.05). All of the discovered motifs share a strong purine preference near the sensed position, and the two SD motifs SD-AA and SD-ME are highly similar (TOMTOM *E*-value: 1.24 *×* 10^*−*7^). The second AAII dwell time motif, DT-AA-2, included a TRA-motif at the sensing region.

In summary, the positional regression revealed sequence and position specificity for the shift in the low quantile values, and the motif discovery revealed sequence logos that were significantly associated with changes in the ionic current. These motifs had purine bases, primarily adenines, at the upper pore positions in AAII and melphalan samples and for all current statistics.

### 3.7 Read interruptions are associated with upstream guanine repeats at the motor protein

Nanopore sequencing occasionally yields incomplete reads due to interruptions in pore translocation [38]. By examining rear barcode qualities (Methods), we observed that AAII and melphalan samples produced more interrupted reads lacking a rear barcode than control samples (Fig. 6A, Suppl. Fig. 1). To investigate whether these interruptions are associated with particular sequence contexts, we performed a motif discovery analysis on interrupted reads. We identified two types of interrupted reads based on the ionic current level at the read end: passing-like reads where the current level at the read end (last 20 measurements) was similar to that of fully-read reads (≥ 50pA), likely resulting from DNA breakage after library preparation, and blocking reads where the level was lower (*<* 50pA), likely due to a bulky adduct obstructing DNA translocation through the pore (Suppl. Fig. 4).

**Figure 6:**
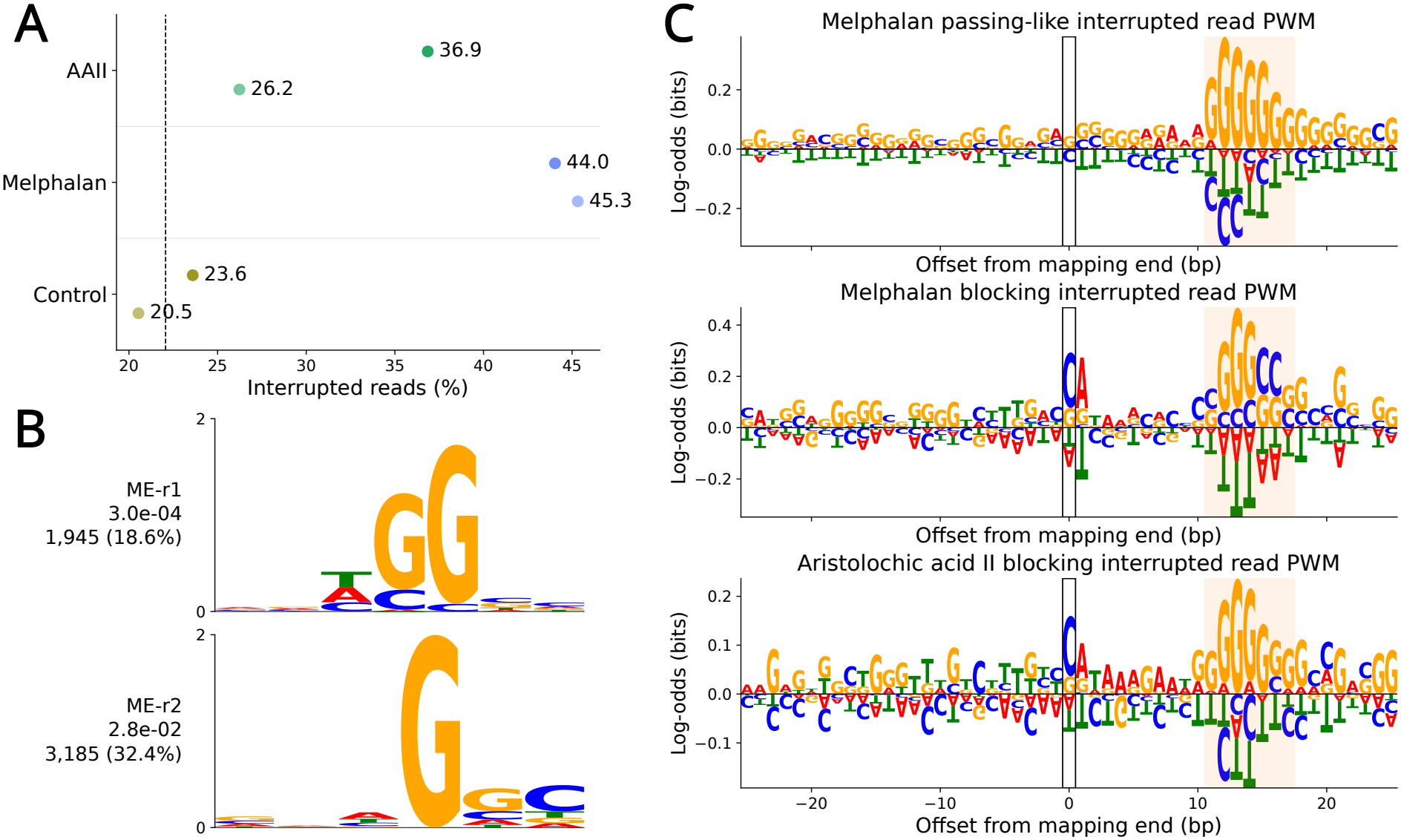
**A:** Fraction of interrupted reads in each sample replicate. The dashed line highlights the average interrupted read fraction in control samples. **B:** A sequence motif associated with read interruptions in blocking melphalan reads at the motor-protein region, when compared against blocking control reads. **C:** Position weight matrices (PWMs) for sequences around the mapping end positions of interrupted reads. The motor-protein region is highlighted in yellow, around +12 bases downstream of the sensing region.

We discovered a guanine-rich motif in the melphalan samples that was associated with the motor-protein position in blocking interrupted reads (*E <* 0.05; Fig. 6B), when compared against control blocking interrupted reads. Additionally, all interrupted reads in AAII and melphalan were enriched for guanines at the motor-protein region, highlighted in yellow in (Fig. 6C), when normalized by the control interrupted read sequences. This led us to believe that both AAII and melphalan monoadducts may cause interruptions in nanopore sequencing. The GGGCC-motif in the motor-protein region in melphalan-blocking reads could suggest a binding position for an interstrand crosslink, which would interrupt sequencing by interacting with the motor protein.

## 4 Discussion

In this study, we investigated how genotoxic, adduct-forming compounds influence nanopore sequencing readouts. To our knowledge, this was the first study to thoroughly examine the effect of DNA adducts in complex DNA contexts (99.96% of all 9-mers) for multiple compounds, whereas many previous studies have focused on specific genomic sites [22, 23, 39] or a specific compound [26]. By applying our novel IonStats framework, we profiled adduct-induced changes across multiple dimensions of nanopore data, including base quality, nanopore current statistics, pore translocation (dwell) times, and read interruptions. This effort revealed a multitude of compound-specific alterations in ionic current properties and sequence contexts, highlighting both shared and distinct patterns of how different adducts perturb DNA translocation through the pore. Our observations establish a foundation for linking nanopore-based adduct detection to exposure-associated mutational processes and signatures [40, 6].

Detecting DNA adducts is a crucial step toward understanding how mutational signatures arise [5, 4, 6]. For instance, signature SBS4 found in lung cancers has been linked to tobacco carcinogens [41]. Site-directed mutagenesis studies demonstrate that the mutational signatures of DNA adducts are directly related to DNA polymerase activity, its DNA insertion and extension mechanism, and the DNA sequence context [42]. To properly assign an etiology to a mutational signature, it is essential to consider DNA adduct types and locations as the key intermediate steps that yield somatic mutations. This, however, requires accurate, site-specific methods for adduct detection: we here showed that nanopore sequencing coupled with sophisticated computational tools for ionic current analysis shows substantial promise.

Our software package IonStats can be easily applied to sets of treated and control samples to evaluate sequencing statistics for treatment effects. In contrast to previous DNA and RNA modification detection tools that can be used to characterize DNA adducts [37, 22, 43], IonStats was developed specifically for DNA adduct detection. It introduces several new readouts on current-level statistics and translocation dynamics, including dwell times and read interruptions. These readouts enable multifaceted characterization of effects caused by DNA adducts. We hope that this tool will encourage further research on profiling effects of environmental exposures with nanopore sequencing beyond metagenomics [44].

Our treatment-specific findings show both potential and challenges. Aristolochic acid II (AAII) altered sequencing quality and ionic current mainly in purine-rich contexts at the nanopore sensing region. In addition, guanine repeats were enriched upstream of the sensing region at the motor protein, associating with translocation interruptions. This finding suggests that the AAII-induced aristolactam II-dA (AL-dA) adducts pass through the nanopore more readily, whereas AL-dG adducts may cause read interruptions by interacting with the motor protein. This is consistent with the larger size of AL-dG adducts compared to AL-dA adducts [8]. Melphalan produced shorter and poorer quality reads than the other treatments, and displayed widespread effects in adenine-rich contexts and evidence of read interruptions in CG-rich contexts. This could indicate melphalan crosslinks, binding to two guanines on opposing strands [45], to cause translocation failure at the motor protein, similarly to AAII.

While IonStats enabled comprehensive profiling of treatment effects, it is not able to identify individual reads harboring adducts. Thus, its sensitivity might be limited when adduct frequencies are low. In addition, we only applied IonStats to *in vitro* samples that are expected to carry relatively large amounts of DNA adducts compared to *in vivo* samples [6]. Further work is required to evaluate the utility of IonStats *in vivo*. Another limitation of our work is that IonStats is not able to quantify the amount of adducts in treated samples directly. A complementary mass spectrometry analysis could be used to estimate the fraction of adducted nucleotides [4].

Taken together, our work establishes a framework for studying DNA adducts with nanopore sequencing and demonstrates how different compounds leave distinct signatures in ionic currents. By developing methods for site-specific adduct detection, future studies may bridge the gap between DNA lesions and mutational signatures, clarifying the causal pathways by which exogenous and endogenous exposures contribute to cancer development.

## Data availability statement

All sequencing data for the adduct experiment have been deposited in the European Nucleotide Archive (ENA) at EMBL-EBI under accession number PRJEB101972. Raw signal data (POD5), reference sequences, and ground truth annotations of the modkit validation dataset were obtained from the Oxford Nanopore Technologies Open Data project, available on AWS at s3://ont-open-data/modbase-validation_2024.10/.

## Acknowledgments

We would like to thank Veera Erkkilä, Pinja Perkkiö, Iina Vuoristo, and Inga-Lill Åberg for technical support, and Harri Kangas for discussions. Additionally, we would like to thank Daniyar Karabayev and Guangzhao Cheng for testing IonStats. We acknowledge CSC – IT Center for Science, Finland, for generous computational resources. Figure 1A was created with BioRender.com.

## Author Contributions

Conceptualization, Y.K., K.P., B.S., and E.P.; Methodology, Y.K., D.P., K.P., and E.P.; Software, Y.K.; Investigation, Y.K., and D.P.; Data analysis, Y.K.; Resources, D.P., N.K.v.K., P.J., L.A.A., B.S., and E.P.; Data curation, Y.K., and K.P.; Writing—original draft, Y.K., D.P., K.P., and E.P.; Writing - Review & Editing, N.K.v.K., P.J., L.A.A., and B.S.; Visualization, Y.K.; Supervision, K.P., B.S., and E.P.; Funding acquisition, B.S., and E.P.

## Funding

This study was supported by the Research Council of Finland (#328890 to EP) and the University of Helsinki Doctoral Programme in Integrative Life Science. Additionally, BS was supported by the Research Council of Norway (187615), Helse Sør-Øst, University of Oslo through the Norwegian Centre for Molecular Biosciences and Medicine (NCMBM) (to Sahu group), Norwegian Cancer Society (274630), South-Eastern Norway Health Authority (HSØ) (2025083), Sigrid Jusélius Foundation, and Finnish Cancer Foundation.

## Notes

https://github.com/ykoski/ionstats/

### Competing Interest Statement

The authors have declared no competing interest.

### Summary of Updates

We added a benchmarking experiment to benchmark IonStats with ELIGOS in a methylation detection task. We revised most of our analyses and figures and excluded the results for cisplatin and mitomycin C due to data quality issues.

## References

[1] Byeong Hwa Yun, Jingshu Guo, Medjda Bellamri, and Robert J. Turesky. DNA adducts: Formation, biological effects, and new biospecimens for mass spectrometric measurements in humans. Mass Spectrometry Reviews, 39(1–2):55–82, 2020.

[2] Aziz Sancar, Laura A. Lindsey-Boltz, Keziban Ünsal-Kaçmaz, and Stuart Linn. Molecular mechanisms of mammalian DNA repair and the DNA damage checkpoints. Annual Review of Biochemistry, 73:39–85, 2004.

[3] Axel Guilbaud, Farzan Ghanegolmohammadi, Yijun Wang, Jiapeng Leng, Alexander Kreymerman, Jacqueline Gamboa Varela, Jessica Garbern, Hannah Elwell, Fang Cao, Elisabeth M. Ricci-Blair, Cui Liang, Seetharamsing Balamkundu, Charles Vidoudez, Michael S. Demott, Kenneth Bedi, Kenneth B. Margulies, David A. Bennett, Abraham A. Palmer, Amanda Barkley-Levenson, Richard T. Lee, and Peter C. Dedon. Discovery adductomics provides a comprehensive portrait of tissue-, age- and sex-specific DNA modifications in rodents and humans. Nucleic Acids Research, 51(20):10829–10845, 2023.

[4] Mu Rong Chao, Yuan Jhe Chang, Marcus S. Cooke, and Chiung Wen Hu. Multi-adductomics: Advancing mass spectrometry techniques for comprehensive exposome characterization. TrAC - Trends in Analytical Chemistry, 180(August), 2024.

[5] Stefano Amente, Giovanni Scala, Barbara Majello, Somaiyeh Azmoun, Helen G. Tempest, Sanjay Premi, and Marcus S. Cooke. Genome-wide mapping of genomic DNA damage: methods and implications. Cellular and Molecular Life Sciences, 78(21–22):6745–6762, 2021.

[6] Gunnar Boysen, Ludmil B. Alexandrov, Raheleh Rahbari, Intawat Nookaew, Dave Ussery, Mu Rong Chao, Chiung Wen Hu, and Marcus S. Cooke. Investigating the origins of the mutational signatures in cancer. Nucleic Acids Research, 53(1), 2025.

[7] Thomas A. Rosenquist and Arthur P. Grollman. Mutational signature of aristolochic acid: Clue to the recognition of a global disease. DNA Repair, 44:205–211, 2016.

[8] Marie Stiborová, Volker M. Arlt, and Heinz H. Schmeiser. DNA adducts formed by aristolochic acid are unique biomarkers of exposure and explain the initiation phase of upper urothelial cancer. International Journal of Molecular Sciences, 18(10), 2017.

[9] Michael Alavanja, John A. Baron, Ross C. Brownson, Patricia A. Buffler, David M. DeMarini, Mirjana V. Djordjevic, Richard Doll, Elizabeth T.H. Fontham, Yu Tang Gao, Nigel Gray, Prakash C. Gupta, Allan Hackshaw, Stephen S. Hecht, Kirsti Husgafvel-Pursiainen, Elena Matos, Richard Peto, David H. Phillips, Jonathan M. Samet, Gary Stoner, Michael J. Thun, Jean Trédaniel, Paolo Vineis, H. Erich Wichmann, Anna H. Wu, and David Zaridze. Tobacco smoke and involuntary smoking. IARC Monographs on the Evaluation of Carcinogenic Risks to Humans, 83:1–1413, 2004.

[10] Thomas W. Kensler, Bill D. Roebuck, Gerald N. Wogan, and John D. Groopman. Aflatoxin: A 50-year Odyssey of mechanistic and translational toxicology. Toxicological Sciences, 120(SUPPL.1):28–48, 2011.

[11] M. Austin Argentieri, Najaf Amin, Alejo J. Nevado-Holgado, William Sproviero, Jennifer A. Collister, Sarai M. Keestra, Midas M. Kuilman, Bigina N.R. Ginos, Mohsen Ghanbari, Aiden Doherty, David J. Hunter, Alexandra Alvergne, and Cornelia M. van Duijn. Integrating the environmental and genetic architectures of aging and mortality. Nature Medicine, 31(3):1016–1025, 2025.

[12] Mengqiu Cao and Xinyu Zhang. DNA Adductomics: A Narrative Review of Its Development, Applications, and Future. Biomolecules, 14(9), 2024.

[13] David K. La and James A. Swenberg. DNA adducts: Biological markers of exposure and potential applications to risk assessment. Mutation Research - Reviews in Genetic Toxicology, 365(1–3):129–146, 1996.

[14] Daniel R. Semlow and Johannes C. Walter. Mechanisms of Vertebrate DNA Interstrand Cross-Link Repair. Annual Review of Biochemistry, 90:107–135, 2021.

[15] Natalia Y. Tretyakova, Arnold Groehler, and Shaofei Ji. DNA-Protein Cross-Links: Formation, Structural Identities, and Biological Outcomes. Accounts of Chemical Research, 48(6):1631–1644, 2015.

[16] Romel P. Dator, Kevin J. Murray, Matthew W. Luedtke, Foster C. Jacobs, Fekadu Kassie, Hai Dang Nguyen, Peter W. Villalta, and Silvia Balbo. Identification of Formaldehyde-Induced DNA-RNA Cross-Links in the A/J Mouse Lung Tumorigenesis Model. Chemical Research in Toxicology, 35(11):2025–2036, 2022.

[17] Gunnar Boysen and Intawat Nookaew. Current and Future Methodology for Quantitation and Site-Specific Mapping the Location of DNA Adducts. Toxics, 10(2), feb 2022.

[18] Lieselot Y. Hemeryck, Anneleen I. Decloedt, Julie Vanden Bussche, Karen P. Geboes, and Lynn Vanhaecke. High resolution mass spectrometry based profiling of diet-related deoxyribonucleic acid adducts. Analytica Chimica Acta, 892:123–131, 2015.

[19] Yuan Jhe Chang, Marcus S. Cooke, Chiung Wen Hu, and Mu Rong Chao. Novel approach to integrated DNA adductomics for the assessment of in vitro and in vivo environmental exposures. Archives of Toxicology, 92(8):2665–2680, 2018.

[20] Mikhail F Denissenko, Sundaresan Venkatachalam, Yu Hua Ma, and Altaf A Wani. Site-specific induction and repair of benzo[a]pyrene diol epoxide DNA damage in human H-ras protooncogene as revealed by restriction cleavage inhibition. Mutation Research - DNA Repair, 363(1):27–42, 1996.

[21] Bo Cao, Xiaolin Wu, Jieliang Zhou, Hang Wu, Lili Liu, Qinghua Zhang, Michael S Demott, Chen Gu, Lianrong Wang, Delin You, and Peter C Dedon. Nick-seq for single-nucleotide resolution genomic maps of DNA modifications and damage. Nucleic Acids Research, 48(12):6715–6725, 2020.

[22] Intawat Nookaew, Gunnar Boysen, Piroon Jenjaroenpun, Hua Du, Pengcheng Wang, Jun Wu, Thidathip Wongsurawat, Sun Hee Moon, En Huang, and Yinsheng Wang. Detection and discrimination of DNA adducts differing in size, regiochemistry, and functional group by nanopore sequencing. Chemical Research in Toxicology, 33(12):2944–2952, ec 2020.

[23] Xinjia Zhao, Yuru Liu, Xiaoyu Chen, Zhuang Mi, Wei Li, Pengye Wang, Xinyan Shan, and Xinghua Lu. Detection and Characterization of Single Cisplatin Adducts on DNA by Nanopore Sequencing. ACS Omega, 6(26):17027–17034, jul 2021.

[24] Yunhao Wang, Yue Zhao, Audrey Bollas, Yuru Wang, and Kin Fai Au. Nanopore sequencing technology, bioinformatics and applications. Nature Biotechnology, 39(11):1348–1365, nov 2021.

[25] Miten Jain, Hugh E. Olsen, Benedict Paten, and Mark Akeson. The Oxford Nanopore MinION: delivery of nanopore sequencing to the genomics community. Genome Biology, 17(1), ec 2016.

[26] Gali Bai, Namrita Dhillon, Colette Felton, Brett Meissner, Brandon Saint-John, Robert Shelansky, Elliot Meyerson, Eva Hrabeta-Robinson, Babak Hodjat, Hinrich Boeger, and Angela N. Brooks. SMAdd-seq: probing chromatin accessibility with small molecule DNA intercalation and nanopore sequencing. Nucleic Acids Research, 53(14):1–12, 2025.

[27] Volker M Arlt, Heinz H Schmeiser, and Gerd P Pfeifer. Sequence-specific detection of aristolochic acid-DNA adducts in the human p53 gene by terminal transferase-dependent PCR The plant extract aristolochic acid (AA), a mixture consisting. Technical Report 1, 2001.

[28] Dalia Mohamed and Michael Linscheid. Separation and identification of trinucleotide-melphalan adducts from enzymatically digested DNA using HPLC-ESI-MS. In Analytical and Bioanalytical Chemistry, volume 392, pages 805–817, nov 2008.

[29] Heng Li. Minimap2: Pairwise alignment for nucleotide sequences. Bioinformatics, 34(18):3094–3100, 2018.

[30] Jurgen F. Nijkamp, Marcel van den Broek, Erwin Datema, Stefan de Kok, Lizanne Bosman, Marijke A. Luttik, Pascale Daran-Lapujade, Wanwipa Vongsangnak, Jens Nielsen, Wilbert H.M. Heijne, Paul Klaassen, Chris J. Paddon, Darren Platt, Peter Kötter, Roeland C. van Ham, Marcel J.T. Reinders, Jack T. Pronk, Dick de Ridder, and Jean Marc Daran. De novo sequencing, assembly and analysis of the genome of the laboratory strain Saccharomyces cerevisiae CEN.PK113-7D, a model for modern industrial biotechnology. Microbial Cell Factories, 11(1):36, 2012.

[31] Sam Kovaka, Paul W. Hook, Katharine M. Jenike, Vikram Shivakumar, Luke B. Morina, Roham Razaghi, Winston Timp, and Michael C. Schatz. Uncalled4 improves nanopore DNA and RNA modification detection via fast and accurate signal alignment. Nature Methods, 22(4):681–691, 2025.

[32] Marcus Stoiber. Modified base best practices and benchmarking. EPI2ME Data Releases, Oxford Nanopore Technologies. https://epi2me.nanoporetech.com/mod-validation-data/, mOctober 2024. Dataset available at s3://ont-open-data/modbase-validation_2024.10/. Accessed: 2026-05-25.

[33] Pauli Virtanen, Ralf Gommers, Travis E. Oliphant, Matt Haberland, Tyler Reddy, David Cournapeau, Evgeni Burovski, Pearu Peterson, Warren Weckesser, Jonathan Bright, Stéfan J. van der Walt, Matthew Brett, Joshua Wilson, K. Jarrod Millman, Nikolay Mayorov, Andrew R.J. Nelson, Eric Jones, Robert Kern, Eric Larson, C. J. Carey, İlhan Polat, Yu Feng, Eric W. Moore, Jake VanderPlas, Denis Laxalde, Josef Perktold, Robert Cimrman, Ian Henriksen, E. A. Quintero, Charles R. Harris, Anne M. Archibald, Antônio H. Ribeiro, Fabian Pedregosa, Paul van Mulbregt, Aditya Vijaykumar, Alessandro Pietro Bardelli, Alex Rothberg, Andreas Hilboll, Andreas Kloeckner, Anthony Scopatz, Antony Lee, Ariel Rokem, C. Nathan Woods, Chad Fulton, Charles Masson, Christian Häggström, Clark Fitzgerald, David A. Nicholson, David R. Hagen, Dmitrii V. Pasechnik, Emanuele Olivetti, Eric Martin, Eric Wieser, Fabrice Silva, Felix Lenders, Florian Wilhelm, G. Young, Gavin A. Price, Gert Ludwig Ingold, Gregory E. Allen, Gregory R. Lee, Hervé Audren Irvin Probst, Jörg P. Dietrich, Jacob Silterra, James T. Webber, Janko Slavič, Joel Nothman, Johannes Buchner, Johannes Kulick, Johannes L. Schönberger, José Vinícius de Miranda Cardoso, Joscha Reimer, Joseph Harrington, Juan Luis Cano Rodríguez, Juan Nunez-Iglesias, Justin Kuczynski, Kevin Tritz, Martin Thoma, Matthew Newville, Matthias Kümmerer, Maximilian Bolingbroke, Michael Tartre, Mikhail Pak, Nathaniel J. Smith, Nikolai Nowaczyk, Nikolay Shebanov, Oleksandr Pavlyk, Per A. Brodtkorb, Perry Lee, Robert T. McGibbon, Roman Feldbauer, Sam Lewis, Sam Tygier, Scott Sievert, Sebastiano Vigna, Stefan Peterson, Surhud More, Tadeusz Pudlik, Takuya Oshima, Thomas J. Pingel, Thomas P. Robitaille, Thomas Spura, Thouis R. Jones, Tim Cera, Tim Leslie, Tiziano Zito, Tom Krauss, Utkarsh Upadhyay, Yaroslav O. Halchenko, and Yoshiki Vázquez-Baeza. SciPy 1.0: fundamental algorithms for scientific computing in Python. Nature Methods, 17(3):261–272, 2020.

[34] Timothy L. Bailey. STREME: accurate and versatile sequence motif discovery. Bioinformatics, 37(18):2834–2840, 2021.

[35] Ammar Tareen and Justin B Kinney. Logomaker: Beautiful sequence logos in Python. Bioinformatics, 36(7):2272–2274, 2020.

[36] Shobhit Gupta, John A. Stamatoyannopoulos, Timothy L. Bailey, and William Stafford Noble. Quantifying similarity between motifs. Genome Biology, 8(2):R24, Feb 2007.

[37] Piroon Jenjaroenpun, Thidathip Wongsurawat, Taylor D Wadley, Trudy M Wassenaar, Jun Liu, Qing Dai, Visanu Wanchai, Nisreen S Akel, Azemat Jamshidi-Parsian, Aime T Franco, Gunnar Boysen, Michael L Jennings, David W Ussery, Chuan He, and Intawat Nookaew. Decoding the epitranscriptional landscape from native RNA sequences. Nucleic Acids Research, 49(2):1–13, 2021.

[38] D. Branton and D.W. Deamer. Nanopore Sequencing: An Introduction. World Scientific Publishing Company, 2019.

[39] Ran Liu, Yezhuang Shen, Jie Mao, Chunzheng Li, Yawei Zhang, Mandong Hu, Yizhe Zhang, Qiuying Han, Weili Gong, Liang Chen, Kun He, Tao Zhou, Weihua Li, and Xianxing Xie. Nanopore sequencing identification of dna aristolochic acid adducts. Chemical Research in Toxicology, 39:2031–2046, 10 2026.

[40] Lauri J. Sipilä, Riku Katainen, Mervi Aavikko, Janne Ravantti, Iikki Donner, Rainer Lehtonen, Ilmo Leivo, Henrik Wolff, Reetta Holmila, Kirsti Husgafvel-Pursiainen, and Lauri A. Aaltonen. Genome-wide somatic mutation analysis of sinonasal adenocarcinoma with and without wood dust exposure. Genes and Environment, 46(1):1–13, 2024.

[41] Ludmil B. Alexandrov, Serena Nik-Zainal, David C. Wedge, Samuel A.J.R. Aparicio, Sam Behjati, Andrew V. Biankin, Graham R. Bignell, Niccolò Bolli Ake Borg, Anne Lise Børresen-Dale, Sandrine Boyault, Birgit Burkhardt, Adam P. Butler, Carlos Caldas, Helen R. Davies, Christine Desmedt, Roland Eils, Jórunn Erla Eyfjörd, John A. Foekens, Mel Greaves, Fumie Hosoda, Barbara Hutter, Tomislav Ilicic, Sandrine Imbeaud, Marcin Imielinsk, Natalie Jäger, David T.W. Jones, David Jonas, Stian Knappskog, Marcel Koo, Sunil R. Lakhani, Carlos López-Otín, Sancha Martin, Nikhil C. Munshi, Hiromi Nakamura, Paul A. Northcott, Marina Pajic, Elli Papaemmanuil, Angelo Paradiso, John V. Pearson, Xose S. Puente, Keiran Raine, Manasa Ramakrishna, Andrea L. Richardson, Julia Richter, Philip Rosenstiel, Matthias Schlesner, Ton N. Schumacher, Paul N. Span, Jon W. Teague, Yasushi Totoki, Andrew N.J. Tutt, Rafael Valdés-Mas, Marit M. Van Buuren, Laura Van’T Veer, Anne Vincent-Salomon, Nicola Waddell, Lucy R. Yates, Jessica Zucman-Rossi, P. Andrew Futreal, Ultan McDermott, Peter Lichter, Matthew Meyerson, Sean M. Grimmond, Reiner Siebert, Elías Campo, Tatsuhiro Shibata, Stefan M. Pfister, Peter J. Campbell, and Michael R. Stratton. Signatures of mutational processes in human cancer. Nature, 500(7463):415–421, 2013.

[42] Robert P. Fuchs and Shingo Fujii. Translesion DNA synthesis and mutagenesis in prokaryotes. Cold Spring Harbor Perspectives in Biology, 5(12):1–22, 2013.

[43] Jared T. Simpson, Rachael E. Workman, P. C. Zuzarte, Matei David, L. J. Dursi, and Winston Timp. Detecting DNA cytosine methylation using nanopore sequencing. Nature Methods, 14(4):407–410, 2017.

[44] Mantas Sereika, Rasmus Hansen Kirkegaard, Søren Michael Karst, Thomas Yssing Michaelsen, Emil Aarre Sørensen, Rasmus Dam Wollenberg, and Mads Albertsen. Oxford Nanopore R10.4 long-read sequencing enables the generation of near-finished bacterial genomes from pure cultures and metagenomes without short-read or reference polishing. Nature Methods, 19(7):823–826, jul 2022.

[45] Lawrence F Povirk and David E Shuker. DNA damage and mutagenesis induced by nitrogen mustards, 1994.

