## Supplementary Information for "Compound-specific DNA adduct profiling with nanopore sequencing and IonStats"

Supplementary material

Yrjö Koski[1,2], Divyesh Patel[2,3], Natalia Kakko von Koch[4], Paula Jouhten[4], Lauri Aaltonen[2,5,6], Kimmo Palin[2,5,6,*], Biswajyoti Sahu[2,3,7,*], and Esa Pitkänen[1,2,6,*]

[1]Institute for Molecular Medicine Finland (FIMM), University of Helsinki, Helsinki, Finland
[2]Applied Tumor Genomics Research Program, Faculty of Medicine, University of Helsinki, Helsinki, Finland
[3]The Norwegian Centre for Molecular Biosciences and Medicine (NCMBM), University of Oslo, Oslo, Norway
[4]Department of Bioproducts and Biosystems, School of Chemical Engineering, Aalto University, Espoo, Finland
[5]Department of Medical and Clinical Genetics, University of Helsinki, Helsinki, Finland
[6]iCAN Digital Precision Cancer Medicine Flagship, Helsinki, Finland
[7]Department of Cancer Genetics, Institute for Cancer Research, Oslo, Norway

### 1 Background

#### 1.1 Nanopore sequencing

Nanopore sequencing is a DNA sequencing technology developed by Oxford Nanopore Technologies (ONT), which provided the first nanopore sequencer, MinION, in 2014 [1]. It can be used to directly sequence native DNA and RNA molecules, and it can provide long sequencing reads, unlike the most widely used NGS technology, Illumina sequencing. The core component of a nanopore sequencer is a nanoscale protein pore, that serves as a biosensor, which is embedded in an electrically resistant membrane. A constant voltage is applied across the membrane, which drives negatively charged single-stranded DNA or RNA fragments to translocate from the negatively charged *cis* side to the positively charged *trans* side in an electrolytic solution. The translocating nucleic acid blocks the flow of ions through the pore on the basis of the sequence that resides inside the nanopore structure, which means that the changes in the measured ionic current correspond to the nucleotide sequence. In order to unwind the double-stranded DNA or RNA-DNA molecules and to control the translocation speed, a motor protein is used to unwind the nucleic acid and to ratchet the molecule through the pore in a step-wise manner.

Since the ionic current is determined by the physical structure of the DNA sequence that resides inside the nanopore, DNA modifications can be identified by analyzing the ionic current. Some of the most well-known DNA modifications, such as 5-methylcytosine (5mC) and 5-hydroxymethylcytosine (5hmC), have been studied extensively with nanopore sequencing, and there are many tools to detect them accurately from nanopore sequencing data. The latest models of the state-of-the-art basecaller dorado support basecalling of variety of modified bases including: 4mC, 5mC, 5mCG, 5hmCG, 5mC, 5hmC, and 6mA. However, there are few publications that focus on the detection of DNA adducts that are less abundant in the genome, and that cover a magnitude of modifications differing in size and the mechanism of formation.

#### 1.2 Adduct-forming compounds

##### 1.2.1 Aristolochic acid

Consumption of *Aristolochia* plants can cause rapidly progressive renal disease, where some patients develop carcinomas in the upper urinary tract. [2] This disease has been previously referred to as Balkan endemic nephropathy (BEN) and Chinese herbs nephropathy (CHN) but these terms have been since replaced with aristolochic acid nephropathy (AAN) due to their common origin. [3] The primary genotoxic compounds in *Aristolochia* plants are the two major forms of aristolochic acid, aristolochic acid I (AAI) and aristolochic acid II (AAII). [4] Enzymatic nitroreduction of AAs generates reactive intermediates that bind to the exocyclic amino groups of dA and dG to form aristolactam (AL)-DNA adducts. [5] AL-dA adducts act as robust biomarkers of AA exposure since they can persist for years in the renal cortex. [2] These AL-dA adducts create primarily A-to-T substitutions during DNA replication. [6] Although both AAI and AAII adducts have been detected in DNA isolated from patients with AAN, AAI is significantly more nephrotoxic than AAII in cell and animal models. [7]

##### 1.2.2 Cisplatin

Cisplatin (cis-diamine-dichloro-platinum) is an antitumor agent that is widely used to treat a variety of solid tumor cancers, such as sarcomas and bone, ovary, testis, neck, and head cancers. [8] Cisplatin activates in the aqueous cytoplasm environment due to the displacement of chloride atoms on cisplatin by water molecules. The resulting highly electrophilic molecule can react with nucleophilic sulfhydryl groups of proteins and with nitrogen atoms on nucleic acids, forming intrastrand and interstrand cross-links on N7-positions on purine residues. [9] When the formation rate of DNA cross-links exceeds the ability of the DNA repair system to counteract them, the cell division process becomes impaired leading ultimately to apoptosis. [9] GpG and ApG intrastrand cross-links have been shown to escape DNA repair [10] which be seen in the mutational signature of cisplatin, where primary substitutions were C >T peaks (CCC >CTC) and four T >A peaks (CT >CA). [11]

##### 1.2.3 Melphalan

Melphalan is an antitumor drug that has been used to treat various types of malignancies, including multiple myeloma, ovarian cancer, breast cancer, and neuroblastoma. [12] It belongs to nitrogen mustards, which undergo intramolecular nucleophilic substitutions to form bioactive aziridinium ions which are capable of alkylating DNA. [13] The primary site for alkylation is the N7-position on guanine, but alkylation of N1, N6, N3, and N7 positions of adenine has also been observed. [14, 15] Melphalan forms both monoadducts and intrastrand and interstrand cross-links. The monoadducts on nitrogen atoms may cause DNA replication inhibition, while the intrastrand cross-links formation is associated with cell death or chromosome loss. The intrastrand lesions may be either promutagenic or lethal. [15, 16] In one study, the monoadducts were primarily observed at G*NN (* indicates the position of the alkylation) and NG*N contexts. The cross-links were less abundant, but they were observed in d(GˆCA)-mel or d(CˆGA)-mel contexts. [17]

##### 1.2.4 Mitomycin C

Mitomycin C (MMC) is an antitumor antibiotic and chemotherapeutic agent that alkylates and cross-links DNA. It is clinically used to treat many types of cancer, including bladder, breast, head and neck, and non-small cell lung cancers. [18] It can alkylate DNA both monofunctionally, resulting in covalently bound monoadducts, and bifunctionally, resulting in interstrand and intrastrand crosslinks. [19] MMC is inert toward nucleotides in its original structure but goes through a reduction reaction *in vivo* which creates a highly reactive bis-electrophile intermediate. [20] Activated MMC interacts primarily with guanines and additional studies have shown the specificity of interstrand cross-link formation at 5'-CpG-3' sequences. [21] Additionally, it was shown that these cross-links cause deletions with a 5'-CpG-3' sequence context prevalent in deleted DNA regions. [22]

#### 1.3 Supplementary methods

Full method descriptions, including cisplatin and mitomycin C treatments.

##### 1.3.1 Cisplatin treatment

Cisplatin DNA adduct was prepared by using the protocol in [23] with modifications. Cisplatin (3 mg, 10 µmol) was activated by incubation with $AgNO_3$ (6 mg, 35 µmol) at 37 °C for 6 hours in 50 µl of water. The activation reaction was conducted in a 1.5 ml Eppendorf tube covered with aluminium foil to prevent light exposure. Cisplatin and $AgNO_3$ mixture was centrifuged at 13,000 rpm for 10 minutes, and the clear supernatant containing activated cisplatin was used for DNA-adduct formation. Cisplatin-DNA adduct was formed by incubating DNA (10 µg) and activated cisplatin in 10 mM HEPES buffer, pH 7.0, at 37 °C for 1 hour in the dark. Cisplatin-treated DNA was purified using Phenol-Chloroform extraction and eluted in 10 mM Tris, pH 8.0.

##### 1.3.2 Mitomycin C treatment

Mitomycin C (MMC) intrastrand crosslink DNA adduct was prepared using the protocol in [24]. Amplified yeast genomic DNA was diluted to 30 µl of 0.1 M PBS, pH 7.4 (P5493, Sigma) (6.54 µg DNA in 30 µl PBS, 60 nmol of DNA). MMC was dissolved in 0.1 M PBS, pH 7.4, to prepare a 100 mM stock solution. 0.24 µmol of mitomycin C (2.4 µl of 0.1 M MMC stock solution was diluted to 27 µl in 0.1 M PBS) and 12 µmol of $Na_2S_2O_4$ (3 µl of 40 mM $Na_2S_2O_4$ in water) were added to amplified yeast genomic DNA and incubated in a PCR tube in a thermoshaker at 40 °C, 400 rpm with open lid in fume hood. MMC-treated DNA was purified by using 1.8x Ampure XP beads and eluted in 20 µl 10 mM Tris-HCl, pH 8.0.

MMC interstrand crosslink DNA adduct was prepared using a protocol similar to intrastrand crosslinking, except the incubation was performed in a closed-lid PCR tube, and a total of 60 µmol of $Na_2S_2O_4$ was added in five portions of 12 µmol at a five-minute interval. MMC-treated DNA was purified by using 1.8x Ampure XP beads and eluted in 20 µl 10 mM Tris-HCl, pH 8.0.

##### 1.3.3 DNA sequencing and data collection

Two replicates for each treatment were prepared for sequencing, with an additional cisplatin sample, where the DNA fragments were selected by size. Approximately 400 ng of DNA was used for each sample to prepare the sequencing library with the Native Barcoding Kit 24 V14 (SQK-NBD114.24, Oxford Nanopore Technologies (ONT)) following the manufacturer's protocol. In total, 13 barcodes were used (two for each treatment, two for controls, and one additional for cisplatin size-selected). The constructed libraries were loaded onto the R10.4.1/FLO-PRO114M flow cell on the PromethION 24 device. The sequencing and data collection were performed with the MinKNOW software v24.02.19, which generated .pod5 files containing the raw ionic current measurements for individual DNA molecules.

The cisplatin treatment heavily damaged the DNA, and we observed a small number of reads for both non-size-selected replicates and the size-selected replicate. Because of this, we excluded cisplatin samples from the analyses. For unknown reasons, the yield for the second MMC intrastrand replicate was very low. As we did not observe any treatment-related effects in any MMC samples, it is likely that the sample treatment was unsuccessful, and we excluded all MMC samples from the analyses.

### 2 Supplementary results

Supplementary Table 1: The choice of two-sample test does not change detection performance.

| | | Kolmogorov-Smirnov | | | Cramér-von Mises | | | Anderson-Darling | | |
|---|---|---|---|---|---|---|---|---|---|---|
| Variable | Mod. | Rec. | Prec. | $F_1$ | Rec. | Prec. | $F_1$ | Rec. | Prec. | $F_1$ |
| Current mean | 5mC | 91.8 | 84.2 | 87.9 | 89.6 | 79.7 | 84.4 | 89.5 | 81.4 | 85.2 |
| | 5hmC | 87.7 | 78.1 | 82.6 | 86.7 | 77.0 | 81.5 | 86.7 | 77.1 | 81.6 |
| | 6mA | 50.6 | 44.7 | 47.4 | 49.8 | 44.6 | 47.0 | 51.4 | 46.1 | 48.6 |
| Current SD | 5mC | 76.2 | 78.3 | 77.2 | 74.2 | 76.6 | 75.4 | 74.2 | 77.1 | 75.6 |
| | 5hmC | 74.8 | 72.4 | 73.6 | 72.1 | 70.6 | 71.3 | 73.8 | 71.9 | 72.8 |
| | 6mA | 63.5 | 69.9 | 66.5 | 60.7 | 69.9 | 65.0 | 60.7 | 69.0 | 64.6 |
| Dwell time | 5mC | 66.2 | 71.1 | 68.6 | 67.2 | 71.2 | 69.1 | 68.0 | 71.5 | 69.7 |
| | 5hmC | 65.0 | 61.4 | 63.2 | 65.2 | 63.0 | 64.1 | 66.2 | 63.5 | 64.8 |
| | 6mA | 45.7 | 50.2 | 47.9 | 45.3 | 49.8 | 47.4 | 45.3 | 50.3 | 47.7 |
| Quality score | 5mC | 33.2 | 48.2 | 39.3 | 33.2 | 47.0 | 38.9 | 33.2 | 46.6 | 38.8 |
| | 5hmC | 36.7 | 50.4 | 42.5 | 32.8 | 45.2 | 38.0 | 33.2 | 46.3 | 38.7 |
| | 6mA | 32.2 | 46.1 | 37.9 | 34.0 | 44.9 | 38.7 | 34.2 | 45.0 | 38.8 |

Recall, precision and $F_1$ (all percentages) for the three two-sample statistics on the modkit validation set of 256 modified sites, averaged over two replicates. All three are evaluated at the single configuration used in the main text (decoy FDR 0.002, peak separation 7, snapping to the candidate base, $\pm 1$ bp tolerance), so only the statistic differs. Cramér-von Mises and Anderson-Darling agree to within 1.6 points of $F_1$ in every cell (mean 0.6), and all three agree to within 4.5 points (mean 1.5) – the same magnitude as the spread between replicates of a single statistic (mean 1.5, maximum 3.9).

Supplementary Table 2: Modification detection without candidate-base snapping.

| | | Calls | | $F_1$ at 1 bp | | | Exact-position recall | |
|---|---|---|---|---|---|---|---|---|
| Variable | Mod. | Free | Snap | Free | Snap | $\Delta F_1$ | Free | Snap |
| Current mean | 5mC | 249 | 279 | 0.895 | 0.879 | +0.017 | 0.586 | 0.803 |
| | 5hmC | 260 | 288 | 0.898 | 0.826 | +0.072 | 0.451 | 0.697 |
| | 6mA | 210 | 290 | 0.330 | 0.474 | −0.144 | 0.055 | 0.406 |
| Current SD | 5mC | 246 | 249 | 0.660 | 0.772 | −0.112 | 0.248 | 0.598 |
| | 5hmC | 259 | 264 | 0.687 | 0.736 | −0.048 | 0.209 | 0.557 |
| | 6mA | 222 | 232 | 0.596 | 0.665 | −0.069 | 0.285 | 0.543 |
| Dwell time | 5mC | 292 | 238 | 0.547 | 0.686 | −0.139 | 0.348 | 0.574 |
| | 5hmC | 333 | 271 | 0.587 | 0.632 | −0.044 | 0.250 | 0.484 |
| | 6mA | 276 | 233 | 0.437 | 0.479 | −0.042 | 0.121 | 0.342 |
| Quality score | 5mC | 704 | 176 | 0.336 | 0.393 | −0.057 | 0.215 | 0.260 |
| | 5hmC | 788 | 186 | 0.355 | 0.425 | −0.070 | 0.213 | 0.289 |
| | 6mA | 804 | 179 | 0.330 | 0.379 | −0.049 | 0.176 | 0.242 |
| ELIGOS | 5mC | 748 | 273 | 0.373 | 0.476 | −0.102 | 0.299 | 0.400 |
| | 5hmC | 925 | 287 | 0.372 | 0.433 | −0.061 | 0.084 | 0.299 |
| | 6mA | 842 | 254 | 0.320 | 0.292 | +0.028 | 0.105 | 0.250 |

Snap-free calling reports peak coordinates as called, while snapping, used for the results in the main text, moves each peak to the nearest base of the modified type within $\max(\text{sep} - 1, 1)$ and discards peaks with no such base in range. $F_1$ is scored against the 256 known sites at the ±1 bp tolerance used throughout, with precision over calls and recall over sites. Snapping is shown at the single configuration used in the main text, decoy FDR 0.002 with peak separation 7, which for IonStats is also the point maximising its mean $F_1$ over all twelve cells. Snap-free has no counterpart to that choice, because its merge window has no analogue in the snapped pipeline, so it is given one operating point per variable, chosen on the mean $F_1$ over the three modifications (FDR, separation, merge window): Current mean 0.002, 9, 9; Current SD 0.002, 7, 7; Dwell time 0.01, 2, 5; Quality score 0.002, 2, 1; ELIGOS 0.002, 2, 1. *Exact-position recall* is the share of the 256 sites carrying a call on the modified base itself, at the same operating points and never re-optimised for that criterion; it isolates localisation from detection and is where snapping earns its place. ELIGOS reports a per-position $p$-value and no localisation of its own, so it was put through this same pipeline: score $-\log_{10} p$, decoy calibration against the same null, the same peak exclusion, truth set and tolerance, at full sequencing depth and 100% modification, averaged over two replicates. ELIGOS is held at the same main-text separation for comparability. Dropping snapping costs ELIGOS as much as it costs IonStats, so the penalty is a property of unsupervised localisation rather than of either tool. These results used the Kolmogorov-Smirnov statistic for IonStats.

Supplementary Table 3: Detection along the dilution series for the current mean with the Kolmogorov-Smirnov statistic, mean of two replicates at ∼2400× coverage, computed on the modkit validation dataset. $F_1$ peaks at 50% rather than 100% modified because recall is already saturated while precision is higher: at 100% the caller makes more calls without finding more true sites, so the additional calls are false.

| Modification | Nominal (%) | Achieved fraction | Recall (%) | Precision (%) | $F_1$ | Calls | Sites found |
|---|---|---|---|---|---|---|---|
| 5hmC | 0 | 0.000 | 0.0 | – | 0.0 | 0 | 0 |
| 5hmC | 1 | 0.010 | 0.0 | – | 0.0 | 0 | 0 |
| 5hmC | 2 | 0.020 | 0.0 | – | 0.0 | 0 | 0 |
| 5hmC | 5 | 0.050 | 0.0 | – | 0.0 | 0 | 0 |
| 5hmC | 10 | 0.100 | 0.8 | 100.0 | 1.6 | 2 | 2 |
| 5hmC | 20 | 0.200 | 61.1 | 90.5 | 73.0 | 173 | 156 |
| 5hmC | 50 | 0.500 | 86.7 | 77.3 | 81.8 | 287 | 222 |
| 5hmC | 100 | 1.000 | 87.3 | 77.9 | 82.3 | 287 | 224 |
| 5mC | 0 | 0.000 | 0.0 | – | 0.0 | 0 | 0 |
| 5mC | 1 | 0.010 | 0.0 | – | 0.0 | 0 | 0 |
| 5mC | 2 | 0.020 | 0.0 | – | 0.0 | 0 | 0 |
| 5mC | 5 | 0.050 | 0.0 | – | 0.0 | 0 | 0 |
| 5mC | 10 | 0.100 | 0.6 | 100.0 | 1.2 | 2 | 2 |
| 5mC | 20 | 0.200 | 52.7 | 93.4 | 67.4 | 144 | 135 |
| 5mC | 50 | 0.500 | 90.8 | 90.1 | 90.5 | 258 | 232 |
| 5mC | 100 | 1.000 | 90.6 | 82.1 | 86.2 | 282 | 232 |
| 6mA | 0 | 0.000 | 0.0 | – | 0.0 | 0 | 0 |
| 6mA | 1 | 0.010 | 0.0 | 0.0 | 0.0 | 1 | 0 |
| 6mA | 2 | 0.020 | 0.0 | 0.0 | 0.0 | 1 | 0 |
| 6mA | 5 | 0.050 | 0.0 | 0.0 | 0.0 | 1 | 0 |
| 6mA | 10 | 0.100 | 0.4 | 29.2 | 0.8 | 4 | 1 |
| 6mA | 20 | 0.200 | 11.9 | 38.4 | 18.2 | 80 | 30 |
| 6mA | 50 | 0.500 | 49.0 | 48.6 | 48.8 | 258 | 126 |
| 6mA | 100 | 1.000 | 49.4 | 44.1 | 46.6 | 287 | 126 |

Supplementary Table 4: Peak-separation sweep on the modkit validation set at full depth, current mean with the Kolmogorov-Smirnov statistic, mean of two replicates. The set of called positions is held fixed, so only localisation varies. The snap window is $\max(sep - 1, 1)$. At the default separation of 2, the snap window is a single base and precision collapses; recall changes comparatively little, which is why $F_1$ rather than recovery is reported throughout. 6mA is the exception, peaking at the smallest separations because its effect is delocalised from the sensed position.

| Modification | Separation | Snap window | Calls | Recall (%) | Precision (%) | $F_1$ |
|---|---|---|---|---|---|---|
| 5hmC | 2 | 1 | 778 | 81.5 | 26.8 | 40.4 |
| 5hmC | 3 | 2 | 640 | 74.4 | 29.8 | 42.5 |
| 5hmC | 5 | 4 | 388 | 72.1 | 47.6 | 57.4 |
| 5hmC | 7 | 6 | 288 | 69.7 | 62.1 | 65.7 |
| 5hmC | 9 | 8 | 263 | 68.0 | 66.2 | 67.0 |
| 5hmC | 11 | 10 | 228 | 63.9 | 71.9 | 67.6 |
| 5mC | 2 | 1 | 715 | 87.7 | 31.4 | 46.2 |
| 5mC | 3 | 2 | 574 | 83.4 | 37.2 | 51.5 |
| 5mC | 5 | 4 | 358 | 80.7 | 57.6 | 67.2 |
| 5mC | 7 | 6 | 279 | 80.3 | 73.7 | 76.8 |
| 5mC | 9 | 8 | 254 | 78.9 | 79.4 | 79.1 |
| 5mC | 11 | 10 | 218 | 73.4 | 86.2 | 79.3 |
| 6mA | 2 | 1 | 702 | 74.4 | 27.1 | 39.8 |
| 6mA | 3 | 2 | 604 | 70.5 | 29.9 | 42.0 |
| 6mA | 5 | 4 | 392 | 55.3 | 36.1 | 43.7 |
| 6mA | 7 | 6 | 290 | 40.6 | 35.9 | 38.1 |
| 6mA | 9 | 8 | 220 | 34.4 | 40.0 | 37.0 |
| 6mA | 11 | 10 | 176 | 29.3 | 42.6 | 34.7 |

Supplementary Table 5: Sequencing statistics per replicate. Read counts are all basecalled reads, before quality filtering. Mean QS is the mean of the per-read mean quality scores. Median 9-mer coverage is the median number of observations contributing to a 9-mer in that replicate.

| Sample | Reads | Mean QS | Median read length (bp) | Median 9-mer coverage |
|---|---|---|---|---|
| Melphalan (r1) | 147,351 | 15.17 | 1,778 | 486 |
| Melphalan (r2) | 137,009 | 15.30 | 1,896 | 480 |
| Aristolochic acid II (r1) | 764,798 | 15.81 | 6,522 | 6,981 |
| Aristolochic acid II (r2) | 1,292,817 | 17.06 | 6,873 | 12,312 |
| Mitomycin C intrastrand (r1) | 1,274,539 | 19.89 | 7,331 | 13,155 |
| Mitomycin C intrastrand (r2) | 34 | 8.76 | 5,556 | 2 |
| Mitomycin C interstrand (r1) | 941,540 | 19.90 | 7,403 | 9,884 |
| Mitomycin C interstrand (r2) | 1,201,957 | 19.96 | 7,450 | 12,652 |
| Control (r1) | 505,227 | 19.54 | 7,000 | 4,868 |
| Control (r2) | 703,900 | 19.59 | 7,102 | 6,860 |
| Cisplatin (r1) | 14,996 | 18.17 | 3,664 | 56 |
| Cisplatin (r2) | 9,353 | 18.29 | 3,630 | 22 |
| Cisplatin (size-selected) | 17,420 | 17.16 | 3,660 | 64 |

Supplementary Table 6: Number of $k$-mers reaching significance after Benjamini–Hochberg correction ($\alpha = 0.05$) in each test, for every replicate compared against a control. The 9-mers column is the number tested. These are significance counts, not modification calls: calling uses the control-versus-control decoy threshold and yields roughly 60% as many positions. The quality score reaches significance at almost every 9-mer in the strongly affected replicates while ranking last in detection performance, which illustrates why significance counts at this sample size measure sensitivity rather than usefulness. Mitomycin C and cisplatin are near zero throughout despite full coverage.

| Sample | Test | 9-mers | Quality score | Current mean | Current SD | Dwell | Adj. dwell |
|---|---|---|---|---|---|---|---|
| Aristolochic acid II (r1) | AD | 262,138 | 261,539 | 206,753 | 107,422 | 98,123 | 22,993 |
| | CVM | 262,138 | 261,437 | 174,343 | 73,731 | 89,328 | 13,792 |
| | KS | 262,138 | 261,185 | 166,652 | 68,249 | 86,072 | 13,230 |
| Aristolochic acid II (r2) | AD | 262,138 | 258,570 | 134,013 | 46,969 | 61,532 | 6,076 |
| | CVM | 262,138 | 258,024 | 109,106 | 32,109 | 57,801 | 4,988 |
| | KS | 262,138 | 256,745 | 98,481 | 28,713 | 54,767 | 4,750 |
| Melphalan (r1) | AD | 262,137 | 212,916 | 191,987 | 7,458 | 17,658 | 7,316 |
| | CVM | 262,137 | 207,340 | 144,843 | 6,229 | 16,081 | 6,722 |
| | KS | 262,137 | 194,412 | 138,276 | 5,611 | 13,232 | 5,305 |
| Melphalan (r2) | AD | 262,138 | 188,821 | 173,416 | 5,692 | 13,218 | 0 |
| | CVM | 262,138 | 180,726 | 121,995 | 4,825 | 12,060 | 4,515 |
| | KS | 262,138 | 163,781 | 115,346 | 4,354 | 9,888 | 3,692 |
| Mitomycin C interstrand (r1) | AD | 262,138 | 0 | 0 | 0 | 0 | 0 |
| | CVM | 262,138 | 94 | 2,299 | 11 | 74 | 56 |
| | KS | 262,138 | 81 | 2,040 | 6 | 43 | 51 |
| Mitomycin C interstrand (r2) | AD | 262,138 | 0 | 0 | 0 | 0 | 0 |
| | CVM | 262,138 | 275 | 2,916 | 21 | 80 | 83 |
| | KS | 262,138 | 230 | 2,571 | 16 | 47 | 50 |
| Mitomycin C intrastrand (r1) | AD | 262,138 | 0 | 0 | 0 | 0 | 0 |
| | CVM | 262,138 | 38 | 1,774 | 8 | 35 | 40 |
| | KS | 262,138 | 15 | 1,530 | 3 | 18 | 29 |
| Cisplatin (r1) | AD | 261,912 | 0 | 0 | 0 | 0 | 0 |
| | CVM | 261,912 | 2,527 | 11,397 | 0 | 0 | 0 |
| | KS | 261,912 | 1,232 | 0 | 0 | 0 | 0 |
| Cisplatin (r2) | AD | 259,425 | 0 | 0 | 0 | 0 | 0 |
| | CVM | 259,425 | 155 | 61,624 | 3 | 0 | 0 |
| | KS | 259,425 | 28 | 2 | 0 | 0 | 0 |

Supplementary Table 7: Two-sample Kolmogorov-Smirnov statistics for read-level mean quality score and read length, each replicate against Control (r1). The statistic is signed so that positive values indicate a distribution shifted to higher values. Every comparison reaches $p < 10^{-16}$, including Control (r2) against Control (r1) at a statistic of 0.01, so the statistic rather than the $p$-value is the quantity of interest.

| Sample | Mean QS | Read length |
|---|---|---|
| Melphalan (r1) | −0.4835 | −0.5667 |
| Melphalan (r2) | −0.4667 | −0.5463 |
| Aristolochic acid II (r1) | −0.5663 | −0.0576 |
| Aristolochic acid II (r2) | −0.3994 | −0.0164 |
| Mitomycin C intrastrand (r1) | 0.0402 | 0.0412 |
| Mitomycin C intrastrand (r2) | −0.6974 | −0.2352 |
| Mitomycin C interstrand (r1) | 0.0406 | 0.0512 |
| Mitomycin C interstrand (r2) | 0.0480 | 0.0569 |
| Control (r2) | 0.0096 | 0.0139 |
| Cisplatin (r1) | −0.1141 | −0.4640 |
| Cisplatin (r2) | −0.0855 | −0.5696 |

Mitomycin C intrastrand (r2) yielded 34 reads and is retained to document that failure; its statistics are not interpretable.

Supplementary Table 8: Pearson correlation of the per-9-mer mean and of the per-9-mer standard deviation of each variable, between each replicate and a control, over the 9-mers covered by both with at least 500 observations. Each treated replicate is compared against one control replicate, the two replicates against different ones. Every comparison reaches $p < 10^{-16}$ at these sample sizes, so the statistic rather than the $p$-value is the quantity of interest.

| | | Per-9-mer mean | | | | Per-9-mer standard deviation | | | |
|---|---|---|---|---|---|---|---|---|---|
| Sample | 9-mers | Mean | SD | Dwell | Adj. dwell | Mean | SD | Dwell | Adj. dwell |
| AAII (r1) | 260,417 | 0.9999 | 0.9968 | 0.7864 | 0.9275 | 0.5608 | 0.9163 | 0.0172 | 0.0403 |
| AAII (r2) | 261,546 | 1.0000 | 0.9985 | 0.9105 | 0.9701 | 0.7453 | 0.9637 | 0.0255 | 0.0500 |
| Melphalan (r1) | 127,884 | 0.9996 | 0.9958 | 0.6237 | 0.7638 | 0.7976 | 0.9744 | 0.0127 | 0.0224 |
| Melphalan (r2) | 126,411 | 0.9996 | 0.9961 | 0.6567 | 0.7917 | 0.8101 | 0.9752 | 0.0147 | 0.0244 |
| MMC inter (r1) | 260,417 | 1.0000 | 0.9991 | 0.9716 | 0.9877 | 0.9816 | 0.9941 | 0.0334 | 0.0830 |
| MMC inter (r2) | 261,546 | 1.0000 | 0.9993 | 0.9785 | 0.9909 | 0.9861 | 0.9955 | 0.0333 | 0.0841 |
| MMC intra (r1) | 260,417 | 1.0000 | 0.9992 | 0.9742 | 0.9888 | 0.9832 | 0.9946 | 0.0346 | 0.0834 |
| MMC intra (r2) | 0 | – | – | – | – | – | – | – | – |
| Cisplatin (r1) | 4,111 | 1.0000 | 0.9974 | 0.9025 | 0.9644 | 0.8840 | 0.9736 | 0.0303 | 0.1186 |
| Cisplatin (r2) | 87 | 1.0000 | 0.9980 | 0.9756 | 0.9167 | 0.7283 | 0.9743 | 0.4419 | −0.0932 |
| Control (r2) | 260,417 | 1.0000 | 0.9990 | 0.9685 | 0.9865 | 0.9797 | 0.9933 | 0.0307 | 0.0800 |

Values for MMC intra (r2), Cisplatin (r1), Cisplatin (r2) rest on too few 9-mers passing the 500-observation filter to be interpretable, and are shown only to document that coverage.

Supplementary Table 9: Signed two-sample Kolmogorov-Smirnov statistics comparing the per-9-mer mean of each variable between each replicate and a control. Positive values indicate a distribution shifted to higher values. Each treated replicate is compared against one control replicate, the two replicates against different ones. Every comparison reaches $p < 10^{-16}$ at these sample sizes, so the statistic rather than the $p$-value is the quantity of interest.

| Sample | 9-mers | Current mean | Current SD | Dwell time | Adjusted dwell time |
|---|---|---|---|---|---|
| Aristolochic acid II (r1) | 260,417 | −0.0033 | 0.0518 | 0.1841 | 0.1111 |
| Aristolochic acid II (r2) | 261,546 | −0.0020 | 0.0304 | 0.1186 | 0.0672 |
| Melphalan (r1) | 127,884 | 0.0031 | 0.0192 | 0.0448 | 0.0310 |
| Melphalan (r2) | 126,411 | 0.0029 | 0.0175 | 0.0483 | 0.0323 |
| Mitomycin C interstrand (r1) | 260,417 | −0.0004 | −0.0008 | 0.0039 | 0.0021 |
| Mitomycin C interstrand (r2) | 261,546 | −0.0005 | −0.0020 | 0.0032 | −0.0012 |
| Mitomycin C intrastrand (r1) | 260,417 | −0.0004 | 0.0010 | 0.0045 | −0.0021 |
| Mitomycin C intrastrand (r2) | 0 | – | – | – | – |
| Cisplatin (r1) | 4,111 | 0.0036 | 0.0265 | 0.0469 | 0.0352 |
| Cisplatin (r2) | 87 | 0.0345 | 0.1034 | 0.1034 | 0.1149 |
| Control (r2) | 260,417 | 0.0003 | −0.0007 | −0.0031 | −0.0016 |

Values for Mitomycin C intrastrand (r2), Cisplatin (r1), Cisplatin (r2) rest on too few 9-mers passing the 500-observation filter to be interpretable, and are shown only to document that coverage.

Supplementary Table 10: Signed Kolmogorov-Smirnov statistics for the per-9-mer standard deviation of each variable. Each treated replicate is compared against one control replicate, the two replicates against different ones. Every comparison reaches $p < 10^{-16}$ at these sample sizes, so the statistic rather than the $p$-value is the quantity of interest.

| Sample | 9-mers | Current mean | Current SD | Dwell time | Adjusted dwell time |
|---|---|---|---|---|---|
| Aristolochic acid II (r1) | 260,417 | 0.6411 | 0.2886 | 0.5751 | 0.5153 |
| Aristolochic acid II (r2) | 261,546 | 0.4952 | 0.1924 | 0.4992 | 0.4361 |
| Melphalan (r1) | 127,884 | 0.4678 | 0.0615 | 0.1370 | 0.1177 |
| Melphalan (r2) | 126,411 | 0.4439 | 0.0577 | 0.1166 | 0.1001 |
| Mitomycin C interstrand (r1) | 260,417 | −0.0127 | 0.0017 | 0.0676 | 0.0469 |
| Mitomycin C interstrand (r2) | 261,546 | −0.0135 | −0.0034 | 0.0572 | 0.0396 |
| Mitomycin C intrastrand (r1) | 260,417 | −0.0072 | 0.0029 | 0.0958 | 0.0669 |
| Mitomycin C intrastrand (r2) | 0 | – | – | – | – |
| Cisplatin (r1) | 4,111 | 0.2296 | 0.0705 | −0.1608 | −0.1741 |
| Cisplatin (r2) | 87 | 0.4023 | 0.1839 | −0.2759 | −0.3333 |
| Control (r2) | 260,417 | −0.0054 | −0.0012 | 0.0294 | 0.0208 |

Values for Mitomycin C intrastrand (r2), Cisplatin (r1), Cisplatin (r2) rest on too few 9-mers passing the 500-observation filter to be interpretable, and are shown only to document that coverage.

Supplementary Table 11: Signed Kolmogorov-Smirnov statistics for the per-9-mer low quantile (1st percentile) of each variable. Cisplatin is omitted: its replicates yield too few observations per 9-mer for quantile-based comparison, as stated in Methods. Each treated replicate is compared against one control replicate, the two replicates against different ones. Every comparison reaches $p < 10^{-16}$ at these sample sizes, so the statistic rather than the $p$-value is the quantity of interest.

| Sample | 9-mers | Current mean | Current SD | Dwell time | Adjusted dwell time |
|---|---|---|---|---|---|
| Aristolochic acid II (r1) | 260,417 | −0.2005 | −0.0386 | −0.0344 | −0.1359 |
| Aristolochic acid II (r2) | 261,546 | −0.0901 | −0.0249 | −0.0206 | −0.0784 |
| Melphalan (r1) | 127,884 | −0.1057 | −0.0151 | −0.0544 | −0.2074 |
| Melphalan (r2) | 126,411 | −0.0995 | −0.0121 | −0.0473 | −0.1885 |
| Mitomycin C interstrand (r1) | 260,417 | 0.0024 | −0.0063 | −0.0042 | −0.0080 |
| Mitomycin C interstrand (r2) | 261,546 | 0.0020 | −0.0044 | −0.0031 | −0.0052 |
| Mitomycin C intrastrand (r1) | 260,417 | 0.0016 | −0.0067 | −0.0057 | −0.0119 |
| Mitomycin C intrastrand (r2) | 0 | – | – | – | – |
| Control (r2) | 260,417 | 0.0014 | −0.0039 | −0.0023 | −0.0043 |

Values for Mitomycin C intrastrand (r2) rest on too few 9-mers passing the 500-observation filter to be interpretable, and are shown only to document that coverage.

Supplementary Table 12: Signed Kolmogorov-Smirnov statistics for the per-9-mer high quantile (99th percentile) of each variable. Each treated replicate is compared against one control replicate, the two replicates against different ones. Every comparison reaches $p < 10^{-16}$ at these sample sizes, so the statistic rather than the $p$-value is the quantity of interest.

| Sample | 9-mers | Current mean | Current SD | Dwell time | Adjusted dwell time |
|---|---|---|---|---|---|
| Aristolochic acid II (r1) | 260,417 | 0.0376 | 0.2361 | 0.2647 | 0.1539 |
| Aristolochic acid II (r2) | 261,546 | 0.0216 | 0.1313 | 0.1760 | 0.0951 |
| Melphalan (r1) | 127,884 | 0.0673 | 0.0433 | 0.1101 | 0.0556 |
| Melphalan (r2) | 126,411 | 0.0616 | 0.0389 | 0.1178 | 0.0577 |
| Mitomycin C interstrand (r1) | 260,417 | −0.0017 | 0.0030 | 0.0226 | 0.0076 |
| Mitomycin C interstrand (r2) | 261,546 | −0.0030 | −0.0027 | 0.0153 | 0.0041 |
| Mitomycin C intrastrand (r1) | 260,417 | −0.0009 | 0.0053 | 0.0302 | 0.0106 |
| Mitomycin C intrastrand (r2) | 0 | – | – | – | – |
| Control (r2) | 260,417 | −0.0007 | 0.0019 | 0.0099 | 0.0033 |

Values for Mitomycin C intrastrand (r2) rest on too few 9-mers passing the 500-observation filter to be interpretable, and are shown only to document that coverage.

Supplementary Table 13: Current-drop enrichment as a function of the threshold $\tau$. The control rate is the fraction of events below $\tau$ in Control (r1). The remaining columns are each replicates's rate divided by it. The enrichment is monotone in $\tau$ while the control-versus-control comparison stays within 5% of unity throughout, so the choice of $\tau$ sets the magnitude but not the conclusion.

| $\tau$ | Control (r1) rate | Aristolochic acid II (r1) fold | Aristolochic acid II (r2) fold | Melphalan (r1) fold | Melphalan (r2) fold | Control (r2) fold |
|---|---|---|---|---|---|---|
| −0.80 | 1.00e-03 | 6.19 | 4.11 | 3.59 | 3.41 | 0.99 |
| −1.00 | 5.00e-04 | 9.15 | 5.89 | 4.37 | 4.14 | 0.98 |
| −1.21 | 3.00e-04 | 12.28 | 7.80 | 5.11 | 4.81 | 0.98 |
| −1.40 | 2.00e-04 | 14.95 | 9.44 | 5.81 | 5.45 | 0.97 |
| −1.60 | 1.00e-04 | 17.35 | 10.92 | 6.59 | 6.13 | 0.96 |
| −1.80 | 1.00e-04 | 20.04 | 12.55 | 7.35 | 6.79 | 0.96 |
| −2.00 | 0.00e+00 | 23.34 | 14.57 | 8.02 | 7.36 | 0.95 |

Supplementary Table 14: Effect of tie-breaking noise on the input to motif discovery: Anderson-Darling statistics on dwell time in the adduct replicates, with and without jitter. Overlap is of the top-ranked $k$-mer sets that STREME is run on; the median change is of the statistic itself.

| Sample | 9-mers | Top 5000 overlap (%) | Top 1000 overlap (%) | Spearman $\rho$ | Median change (%) |
|---|---|---|---|---|---|
| Aristolochic acid II (r1) | 262,138 | 99.6 | 99.6 | 0.9960 | 7.0 |
| Aristolochic acid II (r2) | 262,138 | 99.5 | 99.6 | 0.9930 | 9.4 |
| Melphalan (r1) | 262,137 | 97.8 | 98.5 | 0.9860 | 13.2 |
| Melphalan (r2) | 262,138 | 97.6 | 98.3 | 0.9840 | 13.6 |

Supplementary Table 15: Effect of tie-breaking noise on modification detection for the discrete statistics, on the modkit validation set at full depth. $F_1$ in percent, mean of two replicates. Uniform noise of half-width 0.01 was added to break ties, with the generator seeded at 42.

| Statistic | Modification | Test | $F_1$ (no jitter) | $F_1$ (jitter) | $\Delta F_1$ |
|---|---|---|---|---|---|
| Quality score | 5hmC | AD | 37.34 | 38.68 | +1.33 |
| Quality score | 5hmC | CVM | 36.28 | 38.01 | +1.73 |
| Quality score | 5hmC | KS | 42.03 | 42.48 | +0.45 |
| Quality score | 5mC | AD | 39.86 | 38.77 | −1.09 |
| Quality score | 5mC | CVM | 39.17 | 38.90 | −0.27 |
| Quality score | 5mC | KS | 39.44 | 39.31 | −0.14 |
| Quality score | 6mA | AD | 40.44 | 38.85 | −1.60 |
| Quality score | 6mA | CVM | 37.74 | 38.67 | +0.93 |
| Quality score | 6mA | KS | 38.22 | 37.93 | −0.29 |
| Dwell time | 5hmC | AD | 64.76 | 64.82 | +0.06 |
| Dwell time | 5hmC | CVM | 63.68 | 64.11 | +0.43 |
| Dwell time | 5hmC | KS | 62.64 | 63.19 | +0.55 |
| Dwell time | 5mC | AD | 70.21 | 69.67 | −0.54 |
| Dwell time | 5mC | CVM | 68.95 | 69.15 | +0.20 |
| Dwell time | 5mC | KS | 68.35 | 68.56 | +0.20 |
| Dwell time | 6mA | AD | 48.35 | 47.69 | −0.67 |
| Dwell time | 6mA | CVM | 47.49 | 47.44 | −0.05 |
| Dwell time | 6mA | KS | 47.54 | 47.85 | +0.31 |

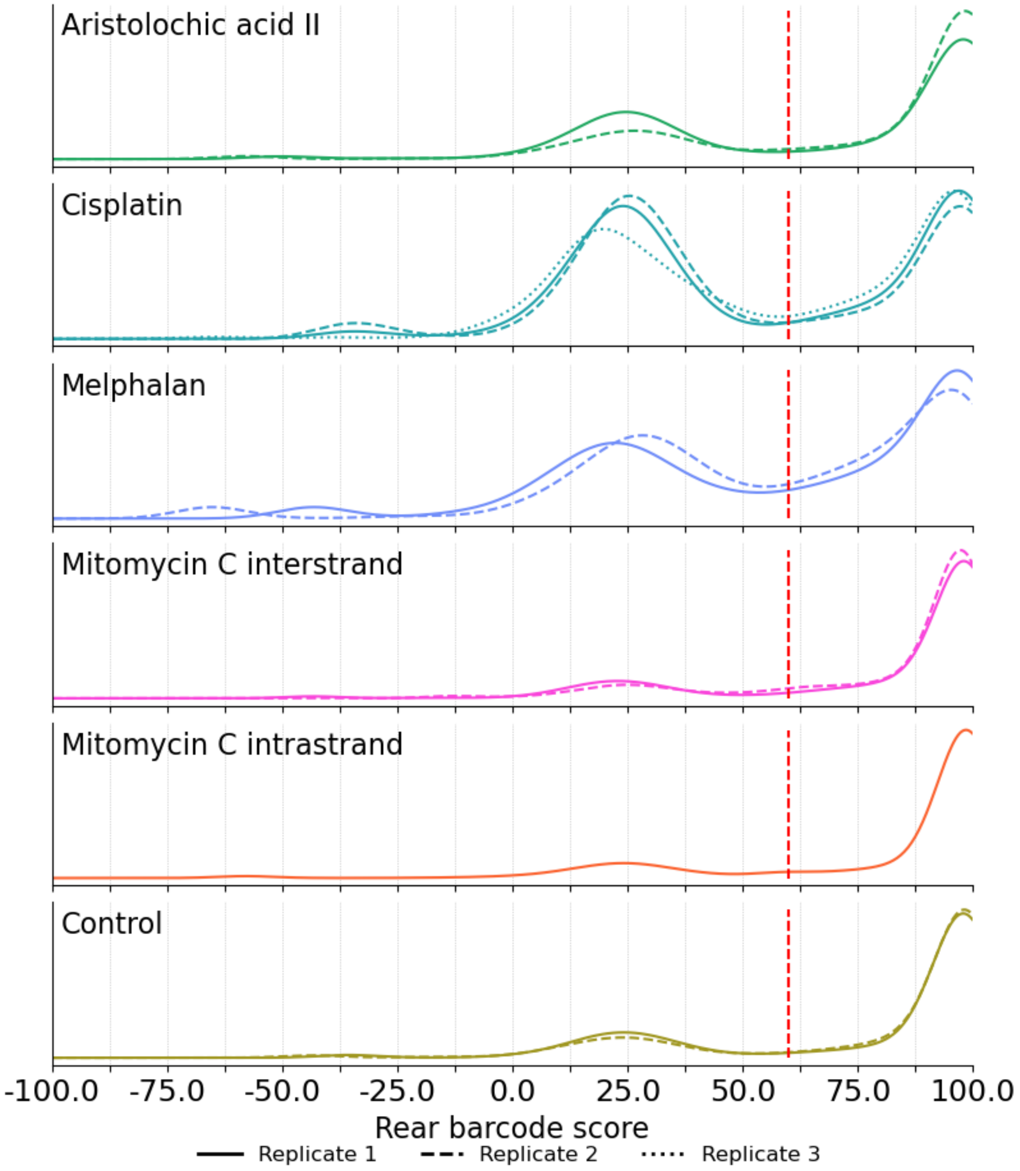


Supplementary Figure 1: Rear barcode score distributions, used to determine interrupted reads, for all treatment groups. The red line signifies the classification threshold (interrupted reads had barcode score $< 60$).

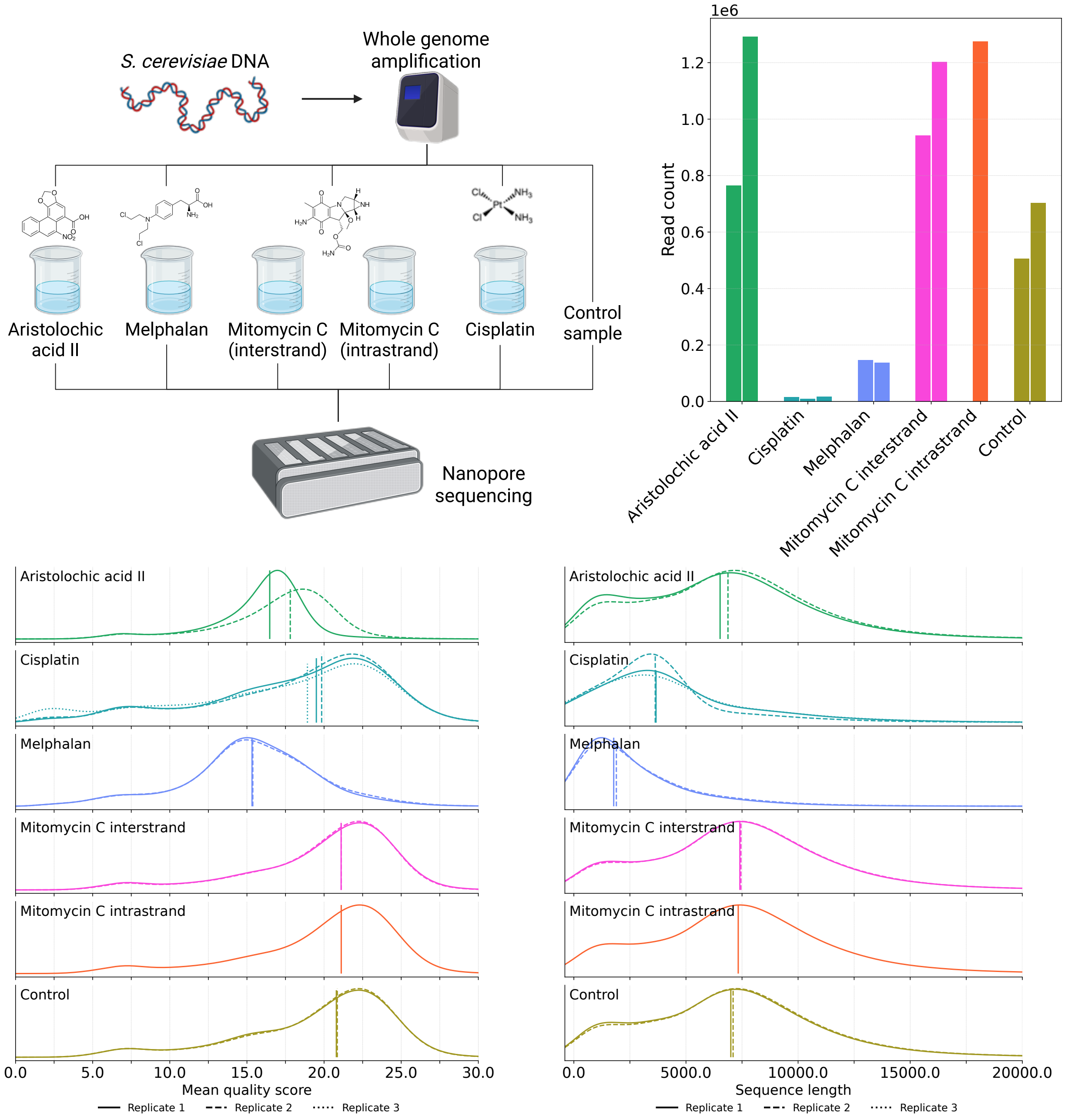


Supplementary Figure 2: Full experimental setup and read counts, mean quality scores, and read lengths for all samples in the dataset.

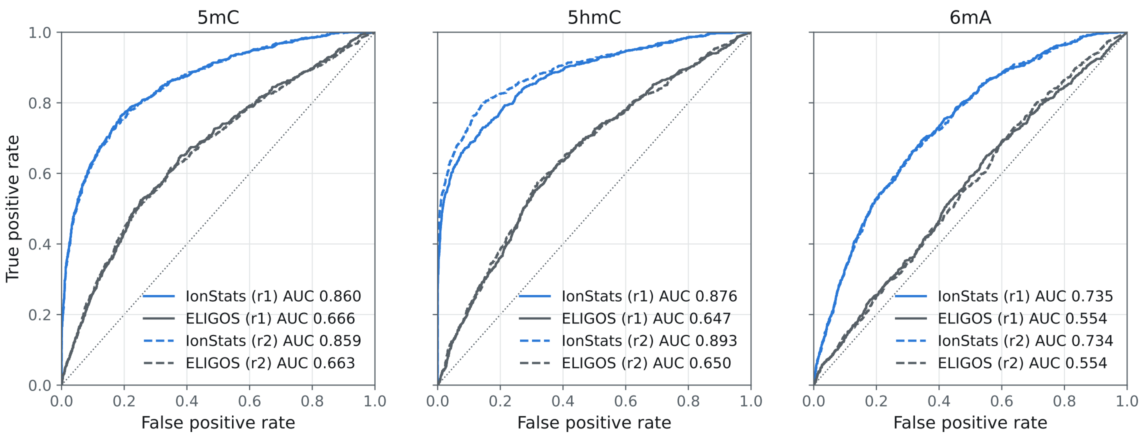


Supplementary Figure 3: ROC-curves of IonStats current mean and ELIGOS on all three methylation types.

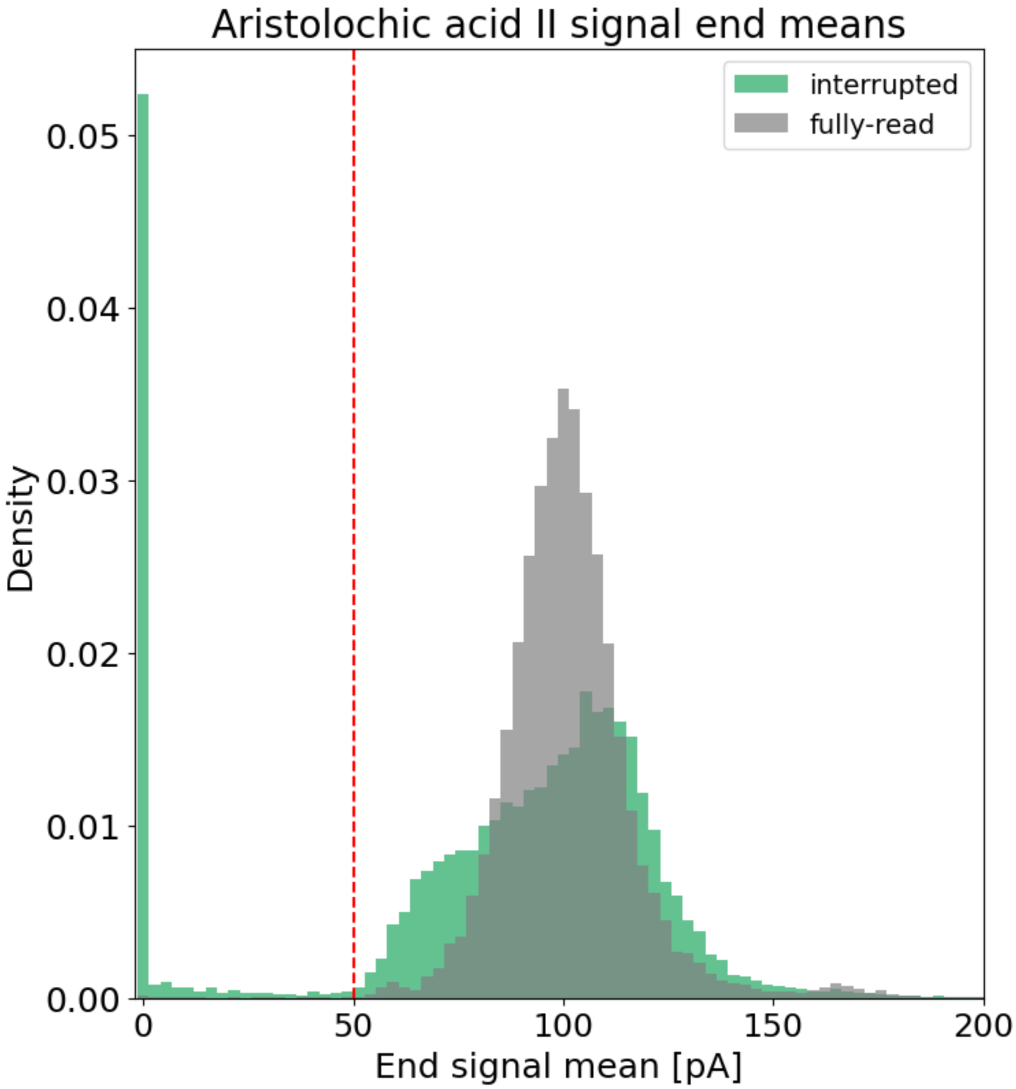


Supplementary Figure 4: Mean signal levels at read ends for interrupted and fully-read reads in pooled AAII replicates. The red line shows the classification threshold for passing-like and blocking reads.

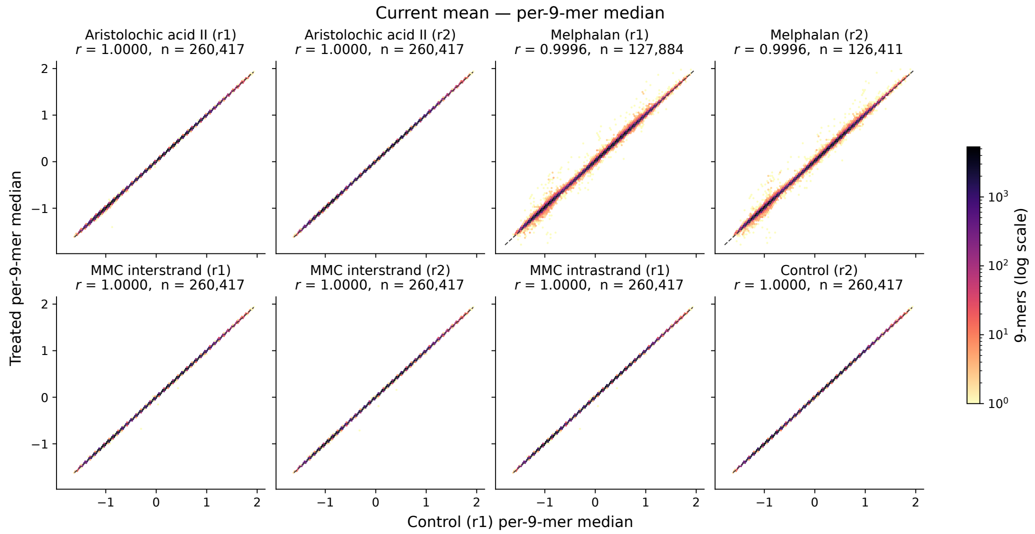


Supplementary Figure 5: Comparison of $k$-mer current mean medians between treated and control samples.

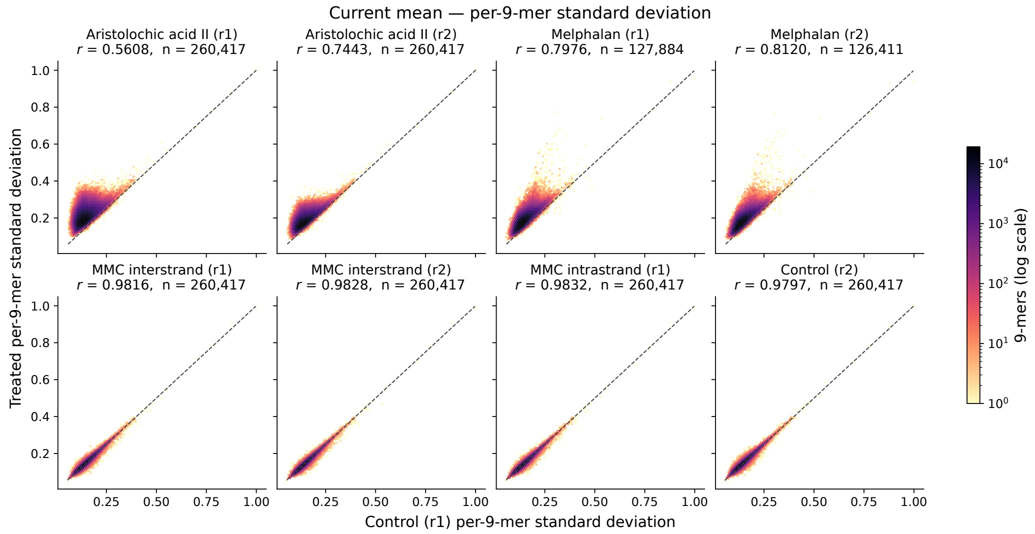


Supplementary Figure 6: Comparison of $k$-mer current mean standard deviations between treated and control samples.

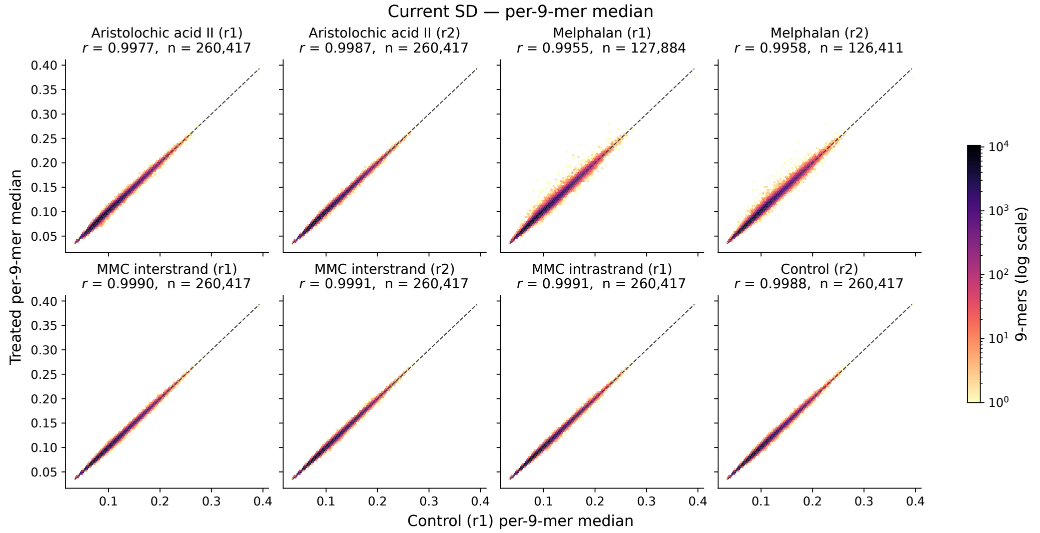


Supplementary Figure 7: Comparison of $k$-mer current SD medians between treated and control samples.

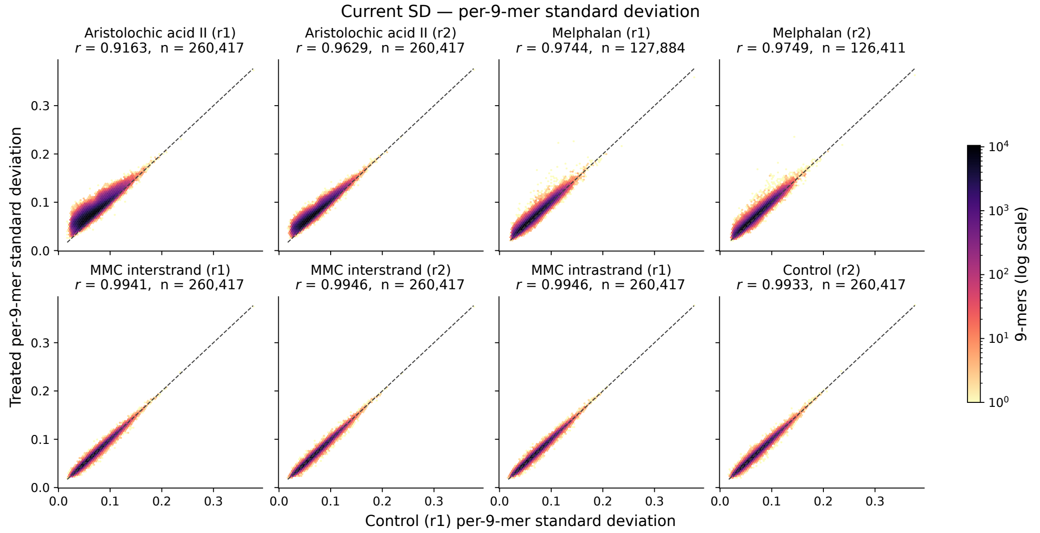


Supplementary Figure 8: Comparison of $k$-mer current SD standard deviations between treated and control samples.

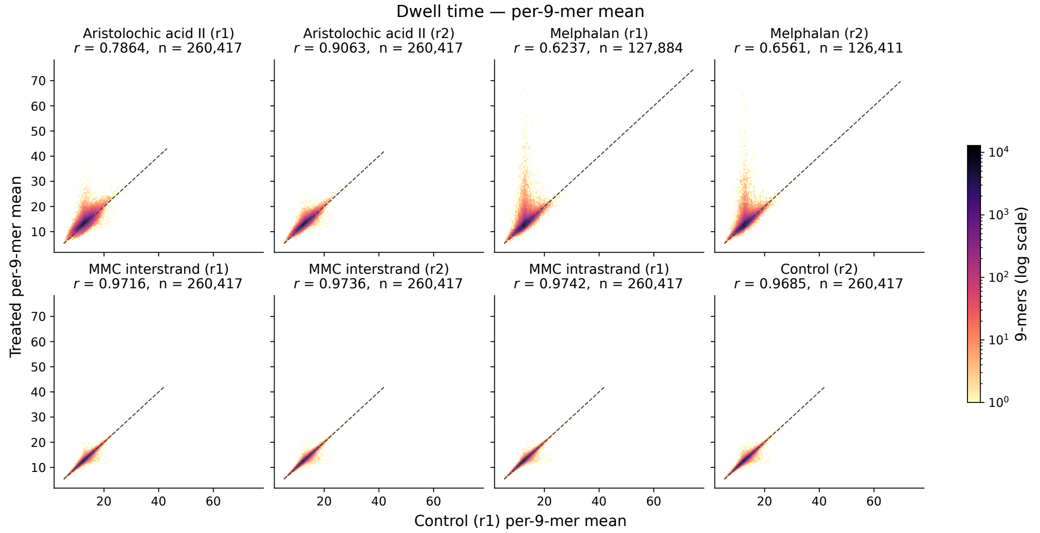


Supplementary Figure 9: Comparison of $k$-mer dwell time means between treated and control samples.

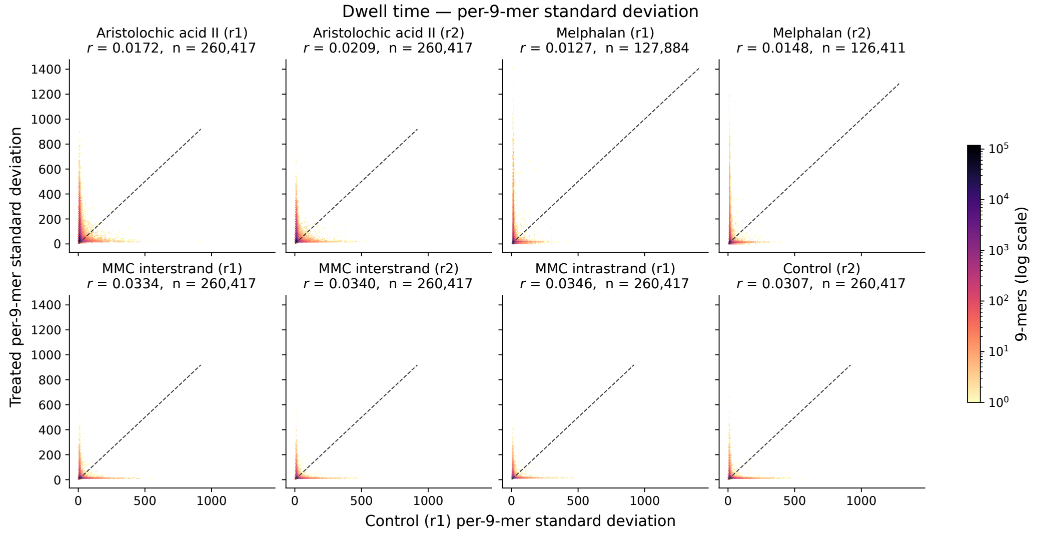


Supplementary Figure 10: Comparison of $k$-mer dwell time standard deviations between treated and control samples.

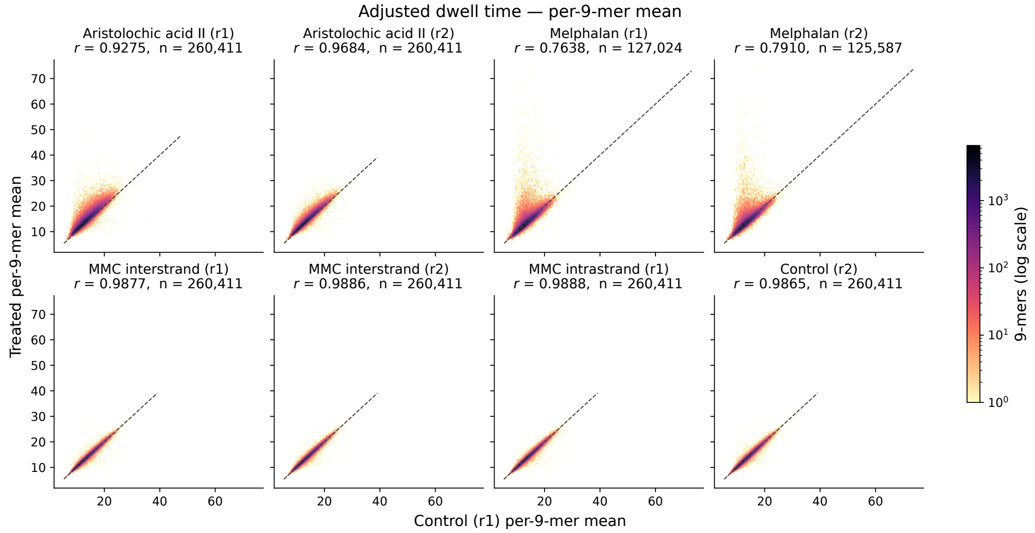


Supplementary Figure 11: Comparison of $k$-mer adjusted dwell time means between treated and control samples.

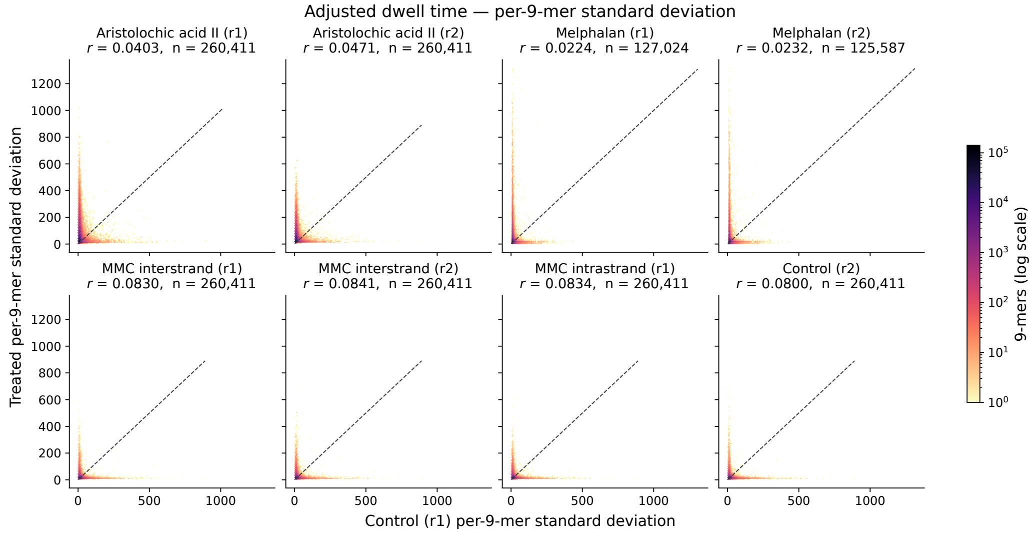


Supplementary Figure 12: Comparison of $k$-mer adjusted dwell time standard deviations between treated and control samples.

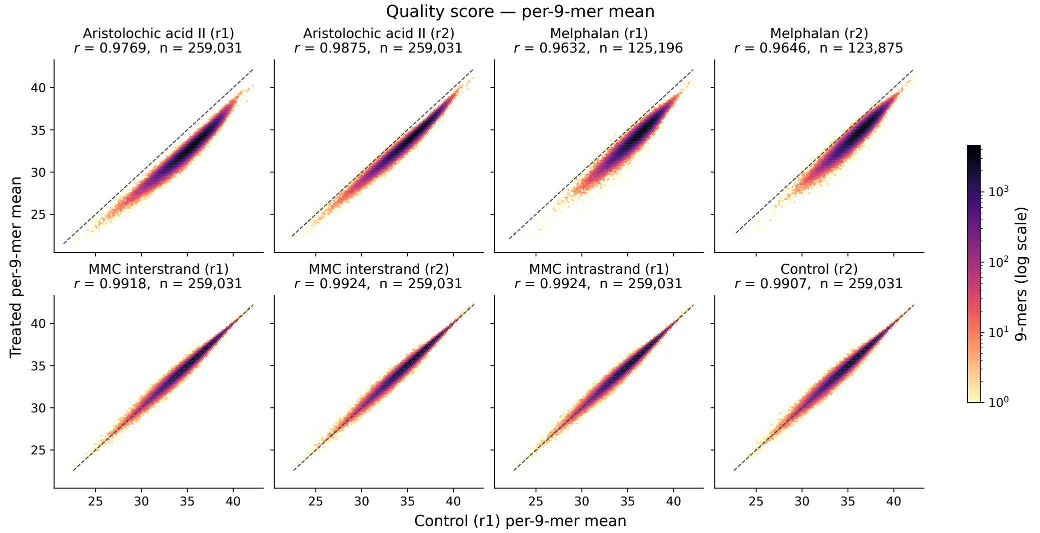


Supplementary Figure 13: Comparison of $k$-mer quality score means between treated and control samples.

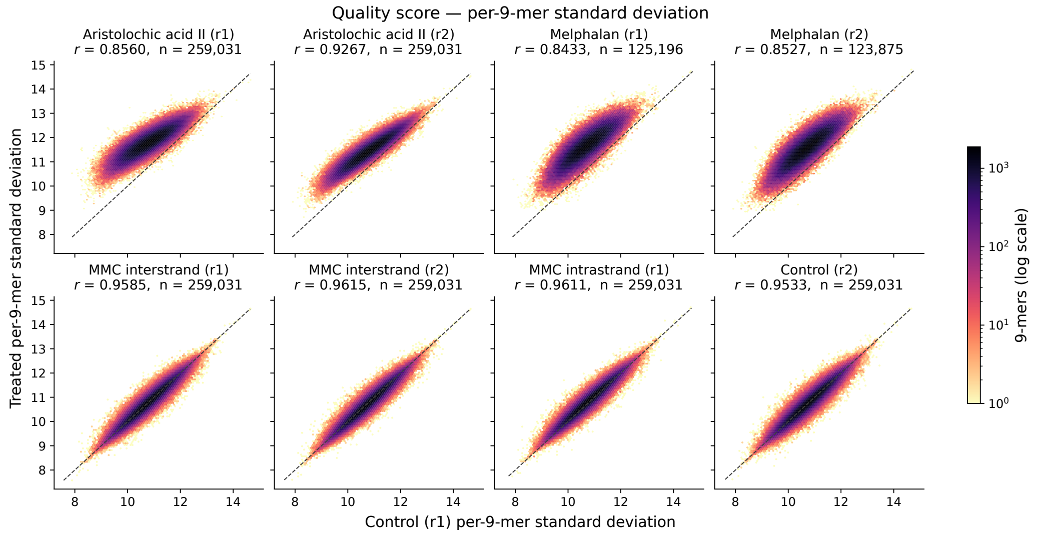


Supplementary Figure 14: Comparison of $k$-mer quality score standard deviations between treated and control samples.

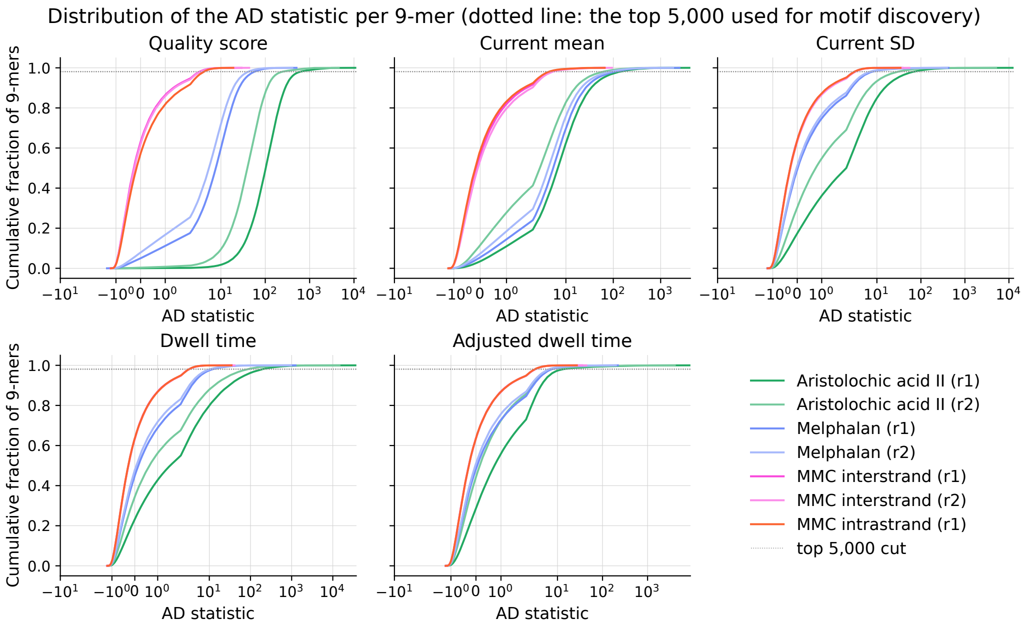


Supplementary Figure 15: AD-test cumulative densities for all treated samples for all studied variables.

### References


[1] Xinjia Zhao, Yuru Liu, Xiaoyu Chen, Zhuang Mi, Wei Li, Pengye Wang, Xinyan Shan, and Xinghua Lu. Detection and Characterization of Single Cisplatin Adducts on DNA by Nanopore Sequencing. *ACS Omega*, 6(26):17027–17034, jul 2021.

[2] Joëlle L. Nortier, Marie-Carmen Muniz Martinzez, Heinz H. Schmeiser, Volker M. Arlt, Christian A. Bieler, Michel Petein, Michel F. Depierreux, and Luc De Pauw. Urothelial carcinoma associated with the intake of a Chinese herb (Aristolochia fangchi). *European Journal of Emergency Medicine*, 8(1):72, 2001.

[3] Marc E De Broe. Chinese herbs nephropathy and Balkan endemic nephropathy : toward a single entity , aristolochic acid nephropathy. *Kidney International*, 81(6):513–515, 2012.

[4] Johanna Michl, Martin J. Ingrouille, Monique S.J. Simmonds, and Michael Heinrich. Naturally occurring aristolochic acid analogues and their toxicities, 2014.

[5] Marie Stiborová, Volker M. Arlt, and Heinz H. Schmeiser. DNA adducts formed by aristolochic acid are unique biomarkers of exposure and explain the initiation phase of upper urothelial cancer. *International Journal of Molecular Sciences*, 18(10), 2017.

[6] Thomas A. Rosenquist and Arthur P. Grollman. Mutational signature of aristolochic acid: Clue to the recognition of a global disease. *DNA Repair*, 44:205–211, 2016.

[7] Noriko Sato, Daisuke Takahashi, Reiko Tsuchiya, Takuya Mukoyama, Shin-ichi Yamagata, Nobunori Satoh, Shiro Ueda, Shih-ming Chen, Makoto Ogawa, Masaaki Yoshida, and Seizo Kondo. Acute nephrotoxicity of aristolochic acids in mice. *Journal of Pharmacy and Pharmacology*, 56(2):221–229, 2004.

[8] Andrea M P Romani. Cisplatin in cancer treatment. *Biochemical Pharmacology*, 206(August):115323, 2022.

[9] Shaloam Dasari and Paul Bernard. Cisplatin in cancer therapy : Molecular mechanisms of action. *European Journal of Pharmacology*, 740:364–378, 2014.

[10] Lloyd R. Kelland. New platinum antitumor complexes. *Critical Reviews in Oncology/Hematology*, 15:191–219, 1993.

[11] Arnoud Boot, Mi Ni Huang, Alvin W.T. Ng, Szu Chi Ho, Jing Quan Lim, Yoshiiku Kawakami, Kazuaki Chayama, Bin Tean Teh, Hidewaki Nakagawa, and Steven G. Rozen. In-depth characterization of the cisplatin mutational signature in human cell lines and in esophageal and liver tumors. *Genome Research*, 28(5):654–665, 2018.

[12] Rakesh Pahwa, Jatin Chhabra, Raj Kumar, and Rakesh Narang. Melphalan: Recent insights on synthetic, analytical and medicinal aspects, 2022.

[13] Lawrence F Povirk and David E Shuker. DNA damage and mutagenesis induced by nitrogen mustards, 1994.

[14] M R Osborne and P. D. Lawley. Alkylation of DNA by melphalan with special reference to adenine derivatives and adenine-guanine cross-linking. *Chemico-Biological Interactions*, 89(1):49–60, 1993.

[15] Shawn Balcome, Soobong Park, R Quirk Dorr, Lucy Hafner, Laura Phillips, and Natalia Tretyakova. Adenine-Containing DNA - DNA Cross-Links of Antitumor Nitrogen Mustards. pages 950–962, 2004.

[16] Scott R Rajski and Robert M Williams. DNA Cross-Linking Agents as Antitumor Drugs. 1998.

[17] Dalia Mohamed and Michael Linscheid. Separation and identification of trinucleotide-melphalan adducts from enzymatically digested DNA using HPLC-ESI-MS. In *Analytical and Bioanalytical Chemistry*, volume 392, pages 805–817, nov 2008.

[18] W T Bradner. Mitomycin C: A clinical update. *Cancer Treatment Reviews*, 27(1):35–50, 2001.

[19] Maria Tomasz and Yolanda Palom. The mitomycin bioreductive antitumor agents: Cross-linking and alkylation of DNA as the molecular basis of their activity, 1997.

[20] Manuel M Paz. Reductive activation of mitomycin C by thiols: Kinetics, mechanism, and biological implications. *Chemical Research in Toxicology*, 22(10):1663–1668, 2009.

[21] Shiv Kumar, Roselyn Lipman, and Maria Tomasz. Recognition of Specific DNA Sequences by Mitomycin C for Alkylation. *Biochemistry*, 31(5):1399–1407, 1992.

[22] Annie S. Tam, Jeffrey S.C. Chu, and Ann M. Rose. Genome-wide mutational signature of the chemotherapeutic agent mitomycin C in Caenorhabditis elegans. *G3: Genes, Genomes, Genetics*, 6(1):133–140, 2016.

[23] Fubo Ma, Shuanghong Yan, Jinyue Zhang, Yu Wang, Liying Wang, Yuqin Wang, Shanyu Zhang, Xiaoyu Du, Panke Zhang, Hong-Yuan Chen, and Shuo Huang. Nanopore Sequencing Accurately Identifies the Cisplatin Adduct on DNA. *ACS Sensors*, 2021.

[24] Manuel M. Paz, Sweta Ladwa, Elise Champeil, Yanfeng Liu, Sara Rockwell, Ernest K. Boamah, Jill Bargonetti, John Callahan, John Roach, and Maria Tomasz. Mapping DNA adducts of mitomycin C and decarbamoyl mitomycin C in cell lines using liquid chromatography/electrospray tandem mass spectrometry. *Chemical Research in Toxicology*, 21(12):2370–2378, dec 2008.
